# *ZNF180* modulates tumor intrinsic immunotherapy resistance in melanoma through driving plasticity

**DOI:** 10.1101/2025.09.16.676573

**Authors:** Won-Min Song, Sree Karani Kondapuram, Xianxiao Zhou, Shu-hsia Chen, Praveen Agrawal

## Abstract

**Background:** Based on our previous study, we have identified *ZNF180*, a zinc finger protein, as a pro-tumorigenic regulator in primary melanoma and a marker for poor prognosis. Herein, we report that *ZNF180*-regulated pathway, hence *ZNF180*-regulome, underlies resistance towards immune checkpoint inhibitions (ICIs).

**Methods:** To investigate regulatory roles of *ZNF180* to confer these immune suppressive phenotypes, we performed *ZNF180* knock-down in melanoma cells *in vitro* with different genetic backgrounds, namely A375 (*BRAF*-mutant) and SKMEL147 (*NRAS*-mutant) cells, and performed RNA- and ATAC-sequencing. We performed integrative analysis of RNA- and ATAC-sequencing data with publicly available sequencing data from ICI-treated cohorts to construct comprehensive model of *ZNF180*-regulome and its impacts on immune microenvironment. Further, we performed *ZNF180* silencing in immune competent Yumm1.7 murine model to confirm the changes in immune microenvironments.

**Results:** *ZNF180*-regulome was predictive of ICI responses in independent bulk sequencing cohorts, and *ZNF180+* tumors persisted after the therapy with immune-suppressive features such as MHC-I loss and CD155 expressions, the primary ligand to TIGIT inhibitory receptor. Further, *ZNF180* silencing revealed its regulations on AP-1 transcription factors to drive melanoma reprogramming towards de-differentiated MITF^low^AXL^high^ cells, an established melanoma subtypes associated with recurrence and ICI resistance. In tandem, we observed that ZNF180+ tumor neighborhood significantly excluded with CD4 T-cells in metastatic tumor, and its silencing in immune competent murine model increased CD4 helper T-cell infiltrations with significant tumor regression *in vivo*.

**Conclusion:** Collectively, these results indicate *ZNF180* is a tumor intrinsic regulator of melanoma plasticity to drive de-differentiated phenotypes with immune-suppressive features including loss of immunogenicity, T-cell inhibitory signals through TIGIT/CD155 checkpoint and exclusion of CD4 helper T-cells. As *ZNF180*-regulome manifests in non-metastatic melanoma in contrast to the current focus of standard-of-care ICI on the metastatic disease, these results establish *ZNF180*-regulome as a biomarker and novel therapeutic avenue for early-stage, non-metastatic melanoma to intervene ICI resistance.

## BACKGROUNDS

T-cell mediated anti-tumor capacities have been successfully translated to standard-of-care via immune-checkpoint inhibition (ICI) therapy, blocking immune-inhibitory signals to PD-1/PD-L1 (anti-PD-1) or CTLA-4/B7 (anti-CTLA-4) co-inhibitory checkpoint axes(1,2). Monotherapies or combinations of these ICIs with BRAF/MEK targeted therapies have collectively improved the 5-year overall survival rates of advanced metastatic diseases from less than 10% to 50%(3). Despite these, many still develop resistance to the therapy eventually, and this represents currently unmet clinical needs.

Currently, most of combination therapies targeting to overcome the ICI monotherapy resistance are focused on advanced metastatic diseases. These efforts include combination of complementary immune checkpoint inhibitions(4): nivolumab (anti-PD-1) with ipilimumab (anti-CTLA4) in NCT02519322 trial (resectable stage III-IV), nivolumab with relatimab (anti-LAG3) in RELATIVITY-047 (unresectable metastatic). Or harbor immunomodulatory effects of genome instability-inducing therapies via increasing neoantigen loads: dacarbazine with nivolumab in NCT00324155 (untreated stage III or dacarbazine-treated stage IV), and radiation therapy with ipilimumab (stage IV)(5,6). Primary ICI resistance accounts for 40-65% of the cases, and is driven by tumor intrinsic factors affecting tumor immunogenicity(7). Although these could manifest in early stages, the current efforts are focused on advanced, metastatic diseases, and the opportunities to detect and intervene them at early stages remain untapped.

Recently, we reported *ZNF180*, a Cys2His2 (C2H2) zinc finger protein, which can recognize DNA base-pairs in a sequence-specific manner, as a multi-functional regulator in primary melanoma(8). While genomic rearrangements on *ZNF180* were associated to DNA repair deficiencies and cellular malignancies(9), high *ZNF180* was predictive of poor overall survival in primary melanoma in The Cancer Genome Atlas (TCGA) (mostly stage II)(8). Further the gene network model suggested that *ZNF180* is an upstream regulator of intral-tumoral DNA repair pathways via regulating *FANCM*, *MSH2* and *ATR*, melanoma oncogenesis via *PIK3CA*, angiogenesis via regulating PAI-I, chromatin remodelling and spliceosome(8). In turn, *ZNF180* silencing showed robust anti-tumor effects in BRAF-mutant (A375) and NRAS-mutant (SKMEL147) cells *in vitro* and *in vivo*, demonstrating as a potent regulator and therapeutic target in melanoma(8). However, the role of *ZNF180* in ICI resistant tumor cells remains elusive.

In this study, we conducted in-depth investigations to study the role of *ZNF180* in conferring ICI resistance in melanoma (**Figure 1A**). First, we collected publicly available single-cell, spatially resolved and bulk sequencing studies to analyse the etiology of *ZNF180*-high tumor subsets in melanoma tumor microenvironment (TME) and evaluated its persistence towards ICI treatment. Then, we conducted *ZNF180* silencing in *BRAF*- and *NRAS*-mutant melanoma cells and performed ATAC- and RNA-sequencing to investigate *ZNF180*-regulatory mechanisms leading to immune evasive phenotypes in melanoma cells. We validated *ZNF180* as a potent target *in vivo* through conducting *ZNF180* silencing in immune competent Yumm 1.7 murine model(10) to confirm tumor regressions and changes in immune infiltrates. Overall, we present *ZNF180* as a novel regulator of ICI resistance and its mechanisms.

**Figure 1.**
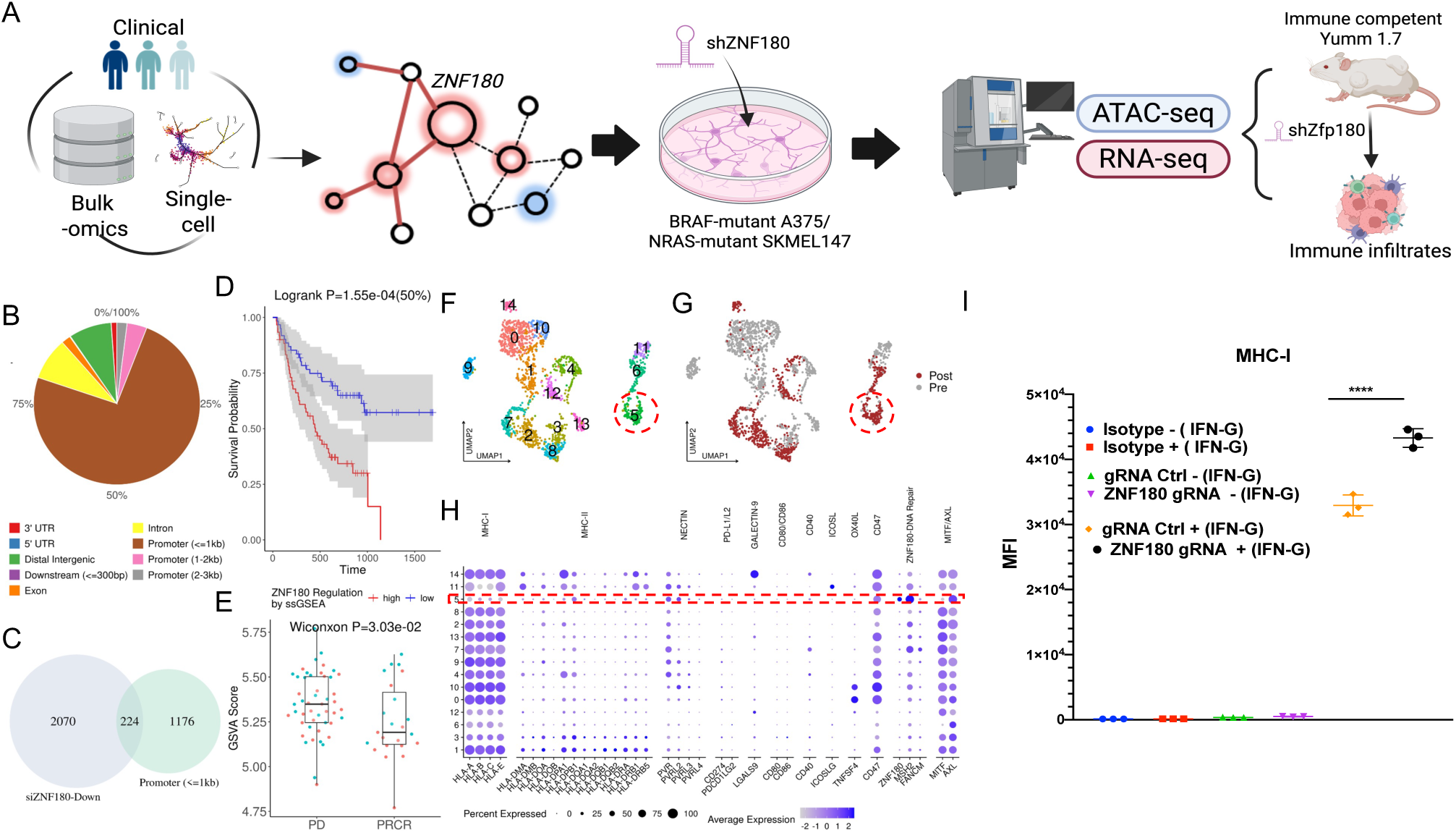
*ZNF180*-high tumors are resistant to anti-PD-1 therapy. **A. Study overview:** Through integrative network modeling of bulk and single-cell sequencing data, *ZNF180* emerged as one of top pro-tumorigenic regulator of melanoma to confer resistances to immune checkpoint inhibition (ICI) therapies. We performed *ZNF180* loss-of-function studies on NRAS- and BRAF-mutant melanoma cells to elucidate tumor-intrinsic changes and *ZNF180*-regulatory network model using ATAC- and RNA-sequencing. The impacts on ICI resistances were further validated through observing changes in key immune modulatory proteins and immune microenvironment changes *in vitro* and immune competent *in vivo* mouse models. **B. *ZNF180* TF binding sites: Annotated peaks from** *ZNF180* TF ChIP-seq from HepG2 cells in ENCODE. **C**. **Putative *ZNF180* regulated pathways in melanoma**: Overlap between down-regulated genes by siZNF180 in SKMEL147 cells (blue) and genes with *ZNF180* binding in promoter sites (≤ 1kb; green). **D. Kaplan-Meier plot of overall survival amongst anti-PD-1 treated metastatic melanoma from Liu *et al.* 2019** ^12^. Patients were stratified by the median of single-sample Gene Set Enrichment Analysis (ssGSEA) score of *ZNF180*-regulated pathways in **C** by GSVA ^25^. **E. Significant activation of ZNF180-regulated pathways in anti-PD-1 non-responders (progressive disease (PR)), compared to the responders (partial/complete response (PRCR)) in Riaz *et al.* 2017** ^26^. Y-axis shows ssGSEA scores of ZNF180-regulated pathways in the Riaz *et al* cohort. **F, G. UMAP plots** of tumor subclusters (**F**), and pre- (grey) and post-anti-PD-1 (brown) status (**G**) in Jerby-Arnon *et al.* 2016 single-cell cohort. **H**. Dotplot of genes (x-axis) in key immune modulatory pathways (labeled at the top). **I.** MHC-I (**J**) surface expression in the parent, and *ZNF180*-knockout B16-F10 (metastatic) cells with or without IFN-γ stimulation for 48 hours and the Mean Fluorescent Intensities (MFI) measured from flow cytometry are presented.

## METHODS

### Experimental Procedures

#### Cell lines and cell culture

Cell line SKmel147 was obtained from Dr. Alan Houghton’s laboratory (MSKCC, New York). The A375 and Yumm1.7 cell line were sourced from American type culture collection. A375 and SKmel147 cells were maintained in DMEM (Corning #10-013-CV), while Yumm1.7 cells were cultured in DMEM-F12 (GIBCO #11320-082). Both media were supplemented with 10% (v/v) fetal bovine serum and 1% (v/v) penicillin/streptomycin. All cell lines were kept in a humidified incubator at 37°C with 5% CO2 and were regularly tested to confirm the absence of mycoplasma contamination.

#### Lentiviral production and transduction of cells with shRNA

Totally, 4 × 10^6^ HEK293T cells were seeded per 10 cm tissue culture plate and incubated overnight at 37 °C and 5% CO_2_. After seeding, HEK293T were co-transfected with bacterial glycerol clones PLKO-puro-shNTC (sigma # SHC016-1EA), ZNF180_shZ1 (shRNA sequence : TGTCCTTGTTGTGCATCAAAG) ZNF180_shZ2 (shRNA sequence : GCTCGACTTATTGTGCATCAA), shZFP180 (shRNA sequence: GAAGTCGTTCATCCAGAGTTA) lentiviral expression constructs (12 μg), viral packaging plasmid (psPAX2, 8 μg), and viral envelope plasmid (VSVG, 4 μg) using Lipofectamine 2000 (Invitrogen # 52887) following manufacturer’s recommendations. Viral supernatant was collected and 0.45 mm filtered at 48 h post-transfection and stored at −80 °C for long-term usage (1-30 days). Cells were transduced with lentivirus in the presence of polybrene (8 μg/ml) and selected with puromycin (1–2 μg/ml) for 4–6 days.

#### RNA extraction, reverse transcription, and qRT-PCR

Total RNA was extracted using miRNeasy QIAGEN mini kit according to the manufacturer’s protocol (QIAGEN #1038703). Totally, 1000 ng of RNA were reverse transcribed using superscript IV Uniprime (Invitrogen #12596025) with random hexamers following the manufacturer’s recommendations. cDNA was diluted with RNase and DNase free water prior to use in a quantitative real-time polymerase chain reaction (qRT-PCR).

GAPDH was used as a housekeeping gene in qRT-PCR. Transcripts were quantified by ABI StepOne Real-Time PCR system (Applied Biosystems #4367659) using Power SYBR Green PCR MasterMix and following 2-step cycling parameters: holding stage, 10 min at 95 °C, and cycling stage 40 cycles of 95 °C for 15 s followed by 60 °C for 1 min followed by melting curve stage. The experiment was performed in three technical replicates.

#### Western blotting

A375 and SkMel147 melanoma cells were lysed in ice-cold RIPA buffer (Thermo Fisher Scientific #89950) with protease inhibitor cocktail (Sigma P8340-ML). After centrifugation at 14,000 rpm (4°C, 20 min), supernatants were collected and protein concentrations determined using the DC protein assay (Bio-Rad #5000114). Protein samples (25 μg) were separated on 4-15% Mini-PROTEAN TGX precast gels (Bio-Rad #4561084) and transferred to PVDF membranes (Millipore IPVH00010). Membranes were rinsed with distilled water and blocked overnight at 4°C in 5% nonfat dry milk in TBST (Tris-buffered saline with 0.05% Tween-20). After brief TBST washing, membranes were incubated overnight at 4°C with primary antibodies diluted 1:1000 in 5% milk-TBST: anti-CD155 (Proteintech #66913-1-1g), anti-PD-L1 (CST #13684), anti-CD58 (Abcam #ab275392), anti-CD112 (CST #95333), anti-MHC-II (CST #68258), anti-B7-H3 (CST #14058), anti-MHC-I (Proteintech #15240-1-AP), anti-Galectin9 (CST #54330), and anti-GAPDH (Proteintech #10494-1-AP). Following three TBST washes, membranes were incubated with HRP-conjugated anti-mouse or anti-rabbit secondary antibodies (1:5000; CST #7074S, Sigma #A9044). After three additional TBST washes, signals were detected using Clarity Western ECL substrate (Bio-Rad #1705062) and visualized on a LICOR Odyssey Fc imaging system.

#### Cell proliferation assay

Transfected cells of Yumm1.7 were plated at a density of 1,000 cells per well in 96-well plates. After allowing cells to adhere overnight, baseline measurements were collected (designated as day 0). Cell viability was subsequently assessed at 24-hour intervals for a total of 4 days using the CellTiter-Glo 2.0 Assay (Promega, Cat# G9242) according to the manufacturer’s protocol. Luminescence was measured at 590 nm using a plate reader. Each experiment included cells transfected with a non-targeting control shRNA (shNTC) as a reference for normalization and experimental control.

#### *In vivo* xenografts

Yumm1.7 cells were infected with shNTC or shZfp180 lentivirus and selected with 2 μg/ml puromycin for 48 h. Cells were, trypsinized, washed with PBS, then suspended in sterile PBS at a concentration of 2 × 10^6^ cells per 150 µl, and maintained on ice until injection. Immediately before injection, cell aliquots were mixed with Matrigel (Corning #354234). Totally, 150 µl of cell/matrigel (1:1) suspensions were injected subcutaneously in the right and left flank of strain C57BL/6J (JAX) (Jackson labs #000664) 4-weeks-old male mice (*n* = 4 per group). When primary tumors were palpable (9 days post-injection), length (*l*) and width (*w*) were measured with calipers 3 times weekly over a period of 14 days. Tumor volume was calculated using the formula (*l* × *w*^2^)/2. Tumor weight was measured at endpoints. Animal experiments were conducted in accordance with guidelines set forth by the Institutional Animal Care and Use Committee (IACUC) of NYU (protocol # 00001527).

#### Tumor dissociation and cell isolation

Subcutaneous tumors were surgically excised from mice and processed immediately. Tumors were washed thoroughly with ice-cold PBS to remove blood contamination. Single-cell suspensions were prepared using the Mouse Tumor Dissociation Kit (Miltenyi Biotec, catalog #130-096-730) according to the manufacturer’s protocol. Tumors were weighed and taken an equal volume of 0.4 g in all conditions. Briefly, tumors were cut into small fragments (2-4 mm) and transferred into gentleMACS C Tubes containing the enzyme mix. Samples were processed using the gentleMACS Dissociator with the appropriate tumor dissociation program, followed by a 30-minute incubation at 37°C under continuous rotation. After the enzymatic digestion, samples were further mechanically dissociated using the gentleMACS Dissociator and filtered through a 70-μm cell strainer to remove any remaining tissue fragments. The resulting cell suspension was washed twice with FACS buffer (PBS containing 2% FBS and 2 mM EDTA). Red blood cells were lysed using RBC lysis buffer (Invitrogen #00-4300-54) for 2 minutes at room temperature, followed by two additional washes with FACS buffer.

#### Sample preparation for flow cytometry

Single-cell suspensions (1×106 cells per sample) were washed with FACS buffer (PBS containing 2% FBS and 2 mM EDTA) and incubated with anti-CD16/CD32 antibody (Fc block, BD Biosciences) for 15 minutes at 4°C to block non-specific binding. For immunophenotyping, cells were stained with two distinct antibody panels. The lymphoid panel included Ghost Dye (Biosciences #13-0863-T100), CD45 (Invitrogen #mcD4517), CD8a (Biolegend #100733), TCRβ (Biolegend #109220), CD4 (Biolegend #100552), and NK1.1 (Biosciences #205941-0025). The myeloid panel consisted of Ghost Dye, CD45, MHCII (Biosciences #107605), CD11c (Biolegend #117335), and LY6C (Biolegend #128035). Antibody staining was performed for 30 minutes at 4°C in the dark, followed by two washes with FACS buffer. Stained cells were resuspended in 300 μl of FACS buffer and maintained at 4°C until flow cytometric analysis.

#### Flow cytometry Analysis

All flow cytometry analysis was performed using a BD FACSymphony A5 SE instrument. Spectral overlap was evaluated with compensation controls, and compensation was calculated automatically. Antibodies for flow cytometry were diluted 1:200, and data analysis was conducted using FlowJo software (FlowJo 10.10.0).

#### ZNF180 knock out in syngeneic B16 cells by CRISPR/Cas9

The parental B16-F0 and metastatic melanoma tumor line B16-F12 were transduced with the Synthego guide RNA of *ZNF180* and Cas-9 Cre mRNA (Synthego.com) using the electroporation. The completely KO cells were established and stimulated with IFN-γ for 48 hours. PD-1 and MHC Class I expression were stained by Flow cytometry, the repeated represented experiment of Mean fluorescent intensities were present.

#### RNA-sequencing library preparation

RNA was isolated from samples using RNeasy® Plus Mini Kit (Qiagen, Hilden, Germany) per the manufacturer’s instructions. Isolated RNA sample quality was assessed by RNA Tapestation (Agilent Technologies Inc., California, USA) and quantified by Qubit 2.0 RNA HS assay (ThermoFisher, Massachusetts, USA). Libraries were constructed with KAPA™ RNA HyperPrep with RiboErase (Roche, Indiana, USA) and performed based on manufacturer’s recommendations. Final library quantity was measured by KAPA SYBR® FAST qPCR and library quality evaluated by TapeStation HSD1000 ScreenTape (Agilent Technologies, CA, USA). The final library size was about 400bp with an insert size of about 250bp. Illumina® 8-nt unique dual-indices were used. Equimolar pooling of libraries was performed based on QC values and sequenced on an Illumina® NovaSeq (Illumina, California, USA) with a read length configuration of 150 PE for 60M PE reads per sample (30M in each direction).

#### ATAC-sequencing data library preparation

ATAC-seq was performed as described previously (Corces et al., 2007). In brief, frozen cells were thawed and washed once with PBS and then resuspended in 500 uL of cold ATAC lysis buffer. The cell number was assessed by Cellometer Auto 2000 (Nexcelom Bioscience, Massachusetts, USA). 50K to 100K nuclei were then centrifugesd at 500g in a pre-chilled centrifuge for 10 min. Supernatant was removed and the nuclei were resuspended in 50 uL tagmentation by pipetting up and down six times. The reactions were incubated at 37 °C for 30 min in a thermomixer with shaking at 300 r.p.m., and then cleaned up by MiniElute reaction clean up kit (Qiagen, Hilden, Germany).

Tagmentated DNA was amplified with barcode primers. Library quality and quantity were assessed with Qubit 2.0 DNA HS Assay (ThermoFisher, Massachusetts, USA), Tapestation High Sensitivity D1000 Assay (Agilent Technologies, California, USA), and QuantStudio ® 5 System (Applied Biosystems, California, USA). Equimolar pooling of libraries was performed based on QC values and sequenced on an Illumina® HiSeq (Illumina, California, USA) with a read length configuration of 150 PE for 200M PE reads (100M in each direction) per sample.

### Bioinformatics Analysis

#### Bulk transcriptome data analysis (Snyder et al. 2014, Riaz *et al.* 2018, Liu et al. 2019)

We sought to obtain the published log-normalized data for bulk transcriptome data. However, it was necessary for some studies to apply additional data processing to obtain the final data. The data processing steps for each cohort are described below,

##### Snyder *et al.* 2014

We accessed log2-transformed, Reads per Kilobase Million (RPKM) normalized data across 21 samples from cBioPortal (https://www.cbioportal.org/study/summary?id=skcm_mskcc_2014)(11).

##### Riaz *et al.* 2018 normalization

Raw gene count matrices were obtained from the published study through Gene Expression Omnibus data portal (GEO accession: GSE91061). The read counts were transformed into Counts Per Million (CPM) values, followed by Trimmed mean of M-values (TMM) scaling with log2-transformation to obtain the log-normalized expressions. These steps were executed through utilizing “*cpm()*” in edgeR R package (v3.38.1).

##### Liu *et al.* 2019 normalization

Transcripts Per Million (TPM)-normalized values from the published study in Supplementary Data 2 were downloaded, and log2-transformed for log-normalization.

##### Survival analysis by *ZNF180* signatures in bulk cohorts

Once the data were collected and pre-processed, we performed single-sample Gene Set Enrichment Analysis (ssGSEA) implemented in “*gsva()*” in GSVA R package (v1.44.1)(12) using *ZNF180* signature derived from intersecting siRNA experiments from our previous study(13) and *ZNF180* TFBS signature from ENCODE(14). Where detailed patient follow-up data with durations and responses were available (i.e. Liu *et al.* 2019 cohort), we performed Cox proportional hazard model to compare prognosis between *ZNF180* signature score-high and -low groups defined by the median score. For Riaz *et al.* 2018 study, we performed t-test between the non-responders (PD) and responders (SD, CR) on the *ZNF180* signature scores to further validate the associations to the responses.

#### Jerby-Arnon *et al.* 2018 data analysis

We downloaded the raw count matrix and cell/sample meta data from Gene Expression Omnibus (GEO) with accession ID, GSE115978. The data were normalized by “*SCTransform()*”(15), then first 20 PCs amongst 2,000 most variable features were used to calculate the UMAP embedding using Seurat R package(16) (v). Then, the tumor cells were subsetted for focused analysis. We used top 10 PCs amongst 2,000 most variable features to perform subclustering on the tumor cells by “*FindClusters()*” function with the default resolution of 0.8 in Seurat R package. This resulted in the 14 subclusters as shown in **Figure 1H**.

#### Biermann *et al.* 2022 spatial transcriptome analysis

We downloaded 16 samples of spatial transcriptome data by SlideSeq-V2 from GSE185386. Using Giotto workflow implemented in Giotto R package (v4.0.8) (17), we filtered out cells with < 100 expressed genes, and genes with < 100 cells with expressions. Then, the expression data were normalized for total library size and scaling by the scale factor of 6,000 using “*normalizeGiotto()*” function. This was followed by adjusting for the number of expressed genes by “*adjustGiottoMatrix()*”. The highly variable genes were identified by “calculateHVF()” with coefficient of variation (COV) z-score > 1.5. The samplewise transcriptomes were integrated by Harmony workflow to synchronize first 20 principal components (PCs) across the samples into a common, harmonized dimension reduction(18). The harmonized PCs were then embedded into two dimensional space by UMAP(19).

The cell types on each data point was inferred using InstaPrism workflow(20) by utilizing Jerby-Arnon *et al.* 2018 scRNA-seq as the reference. While InstaPrism provides the normalized abundances of cell types present at each spatial data point, we determined the cell types present at these data points, whose abundance exceeds > 1/3 by leveraging that the resolution of Slide-seq V2 allows upto 3 cells at each data point. Knowing that *ZNF180* is expressed only in the tumors(13), this further facilitated identification of *ZNF180*+ tumors by pinpointing data points with tumors and *ZNF180* expressions jointly.

Anchored on the *ZNF180*+ tumors, we performed enrichment analysis of different cell types in ZNF180+ tumor neighborhoods. Per data point identified as ZNF180+ tumor, the neighborhood was defined as k nearest neighboring (kNN) data points where we used kNN = 50. Then, the enrichment or depletion of each cell type in the neighborhoods was evaluated by Fisher’s Exact Test (FET)(21) with odds ration (OR) < 1 for depletion, and OR > 1 for enrichment with FDR-adjusted FET p-value < 0.05. We have evaluated the enrichments for kNN=10, 20, 30 and 40 to ensure the robustness of the observed enrichments or depletions.

#### RNA-sequencing data processing

High-throughput RNA-sequencing data is subjected to adapter removal with cutadapt and base quality trimming to remove 3′ read sequences if more than 20 bases with Q<20 were present. The reads are mapped to GRCh38 by STAR aligner^25^, the gene counts for fragments mapped into exonic regions are called by featureCounts^26^, then are normalized by Counts Per Million (CPM), followed by Trimmed Mean of M (TMM) scaling normalization^27^. For each cell line, we performed *DESeq2* analysis^18^ with FDR < 0.05 to identify differentially expressed genes between *ZNF180* silenced cells by two shRNAs (shZ1, shZ3) and non-transfected controls (NTC) with two biological replicates per shRNA per condition (shZ1, shZ3 and NTC)

#### ATAC-sequencing data processing

Reads were mapped to reference genome hg38 using *STAR* aligner(22). Peaks were called for each sample using the Model-based Analysis of ChIP-seq (MACS)(23) (https://github.com/taoliu/MACS/). Quality control (QC) was performed on the samples, such as the percentage of reads in peaks, the number of detected peaks and outlier analysis based on principal component analysis (PCA). Peaks overlap with the “Blacklist” regions by ENCODE(14) were removed. Peaks from all samples were integrated into a single list by union operations. Next, peaks were annotated to genome regions and classified into promoters, exons, introns, 5’UTRs, 3’UTRs and intergenic regions. Peaks overlap with promoters and gene bodies were linked to genes. Enrichment analysis of peaks and genomic regions (such as H3K4me1 peaks and transcription factor binding sites (TFBS)) were performed using the R package *LOLA*(24), in which genomic regions were collected from the ENCODE(14), CODEX(25), Cistrome(26) and ROADMAP(27) databases. For the gene-linked peaks, read counts will be called using featureCounts function from the *Rsubread* package(28) to generate a sample-by-peak read count matrix. Scaling normalization were performed on the matrix using trimmed mean of M-values method(29). Differential accessible peaks (DAPs) were identified between treatment conditions using the R package *limma*(30) with FDR < 0.05. Gene ontology enrichment analysis were performed on the genes linked to DAPs to show the functions of the DAPs.

We integrated the functional DAPs in promoter regions across the cell lines at gene level. For each gene, we queried DAPs in promoter regions that are the nearest to the TSS, and combined the p-values by ACAT(31).

#### Motif enrichment analysis

We utilized Hmm-based IdeNtification of Transcription factor footprints (HINT)(32) from regulatory genomics toolbox (RGT) to perform motif enrichment analysis in the DAPs with FDR < 0.05. The enrichment fold change (EFC) statistics from motif enrichment analysis in A375 and SKMEL147 cells were integrated to identify commonly enriched TF motifs in the DAPs (**Figure 3E**).

#### STRING PPI analysis of ZNF180-regulome

We downloaded the list of protein-protein interactions curated in STRING database (v10.5), extracted the high confidence PPIs (confidence score > 900) and overlapped the consensus DEGs and DAPs from A375 and SKMEL147 cells to identify ZNF180-regulated PPI network, hence *ZNF180* regulome. We further searched for densely connected subnetworks by optimizing the network modularity(33) by walktrap algorithm(34) implemented in cluster_walktrap() function *igraph* R package (v2.0.3). This yielded 9 subnetworks as shown in **Figure 4A**.

## RESULTS

### *ZNF180*-regulated pathways in melanoma cells are predictive of anti-PD-1 responses and modulate MHC-I changes in murine melanoma

Firstly, we verified transcriptional regulations by *ZNF180* as TF in melanoma cells. Using *ZNF180* binding peaks from TF ChIP-seq in HepG2 cells in ENCODE(35) (**Figure 1B**), we confirmed significant overlaps of the promoter site bindings within 1kb of Transcription Starting Site (TSS) with the down-regulated genes upon *ZNF180* silencing in NRAS-mutant SKMEL147 cells from our previous study(8) with Fisher’s Exact Test (FET) FDR=2.08E-7 (**Figure 1C**).

Utilizing the overlap genes between these two signatures as a putative *ZNF180*-regulated pathway in melanoma, we evaluated whether the activation of the *ZNF180*-regulated pathway is associated with ICI outcomes. We observed that stronger activations of the signature were significantly associated with poor overall survival in the bulk transcriptome of 144 anti-PD-1 treated metastatic melanoma from Liu *et al.* 2019(36) (Logrank p = 1.55E-4; **Figure 1D**; see **METHODS**). Further, from an independent cohort of 68 patients treated with PD-1 blockade in Riaz *et al.* 2018(37), we observed significantly stronger activations of *ZNF180*-regulated pathway in non-responders with progressive disease (PD), compared to the responders with partial response (PR) or complete remission (CR) (Wilcoxon p = 3.03E-2; **Figure 1E**; see **METHODS**).

Further, *ZNF180*-high tumors exhibited altered immune modulatory signals. We performed tumor subclustering on scRNA-seq of 33 melanoma tumors from pre- and post-anti-PD-1 therapy from Jerby-Arnon *et al.* 2018(38), (**Figure 1F, G**; see METHODS), and identified subcluster 5 as the *ZNF180*-high tumor. Subcluster 5 showed strong suppression MHC-I and high *MITF/AXL* ratio through down-regulation of *MITF* and up-regulation of *AXL* (**Figure 1H**), a hallmark of the de-differentiated melanoma and resistance to anti-PD-1 therapy(39).

We validated *ZNF180*’s regulatory role on MHC-I *in vitro*. We conducted CRISPR/Cas9 to knock out Zfp180, murine ortholog to human *ZNF180,* in syngeneic melanoma B16-F16 tumor lines (see **METHODS**). The data show that guide RNA (gRNA)-mediated knockout of *Zfp180* with IFN-γ stimulation has significantly increased MHC-I expressions compared to gRNA control in B16-F16 tumor cells (**Figure 1I, Supplemental Figure 1**). These suggest *ZNF180* negatively influences T-cell recognition and ICI efficacy.

Overall, these data support *ZNF180*’s role as a tumor-intrinsic regulator of ICI resistance in melanoma and necessitates further understanding its regulatory mechanisms to explore the potential therapeutic avenues.

### *ZNF180* knockdown robustly silences oncogenic and immunosuppressive pathways

To study the regulatory elements by *ZNF180* in tumor cells with different genetic backgrounds, we performed *ZNF180* loss-of-function study by short hairpin RNAs (shRNAs) on *BRAF*-mutant A375 and *NRAS*-mutant SKMEL147 cells. Per cell line, *ZNF180* expressions were knocked down (KD) by two shRNAs (shZ1 and shZ3) for robustness against the off-target effects, and two biological replicates were produced per group (NTC, shZ1 and shZ3) (see **METHODS** for **RNA-sequencing data processing**). We confirmed that the shRNAs effectively silenced *ZNF180* expressions in these cells by qPCR (**Supplemental Figure 2**).

Then, we performed RNA sequencing on these cells (**Supplemental Figure 3**) to identify differentially expressed genes (DEGs) by *ZNF180* KD (**Figure 2A**; **Supplemental Table 1A**; see **METHODS**). Many DEGs were shared in both cells lines, including 1,860 up-regulated (UP) and 1,094 down-regulated (DN) genes shared with FET p-value < 1E-320 (**Figure 2B**). Log2(fold changes) from the two cell lines also showed concordances with significant correlations (Pearson correlation = 0.242, P < 2.2E-16; **Figure 2C**). The commonly up- or down-regulated genes in both cell lines served as the consensus DEG signatures (**Figure 2C**). Of those, we observed the down-regulated DEGs were significantly enriched for transcriptionally controlled genes by *ZNF180*. Utilizing *ZNF180* transcription factor binding sites (TFBS) curated by ENCODE(14) (q-value < 0.05), promoter sites of 158 genes in the consensus down-regulated signature included the *ZNF180* TFBS (**Figure 2D**).

**Figure 2.**
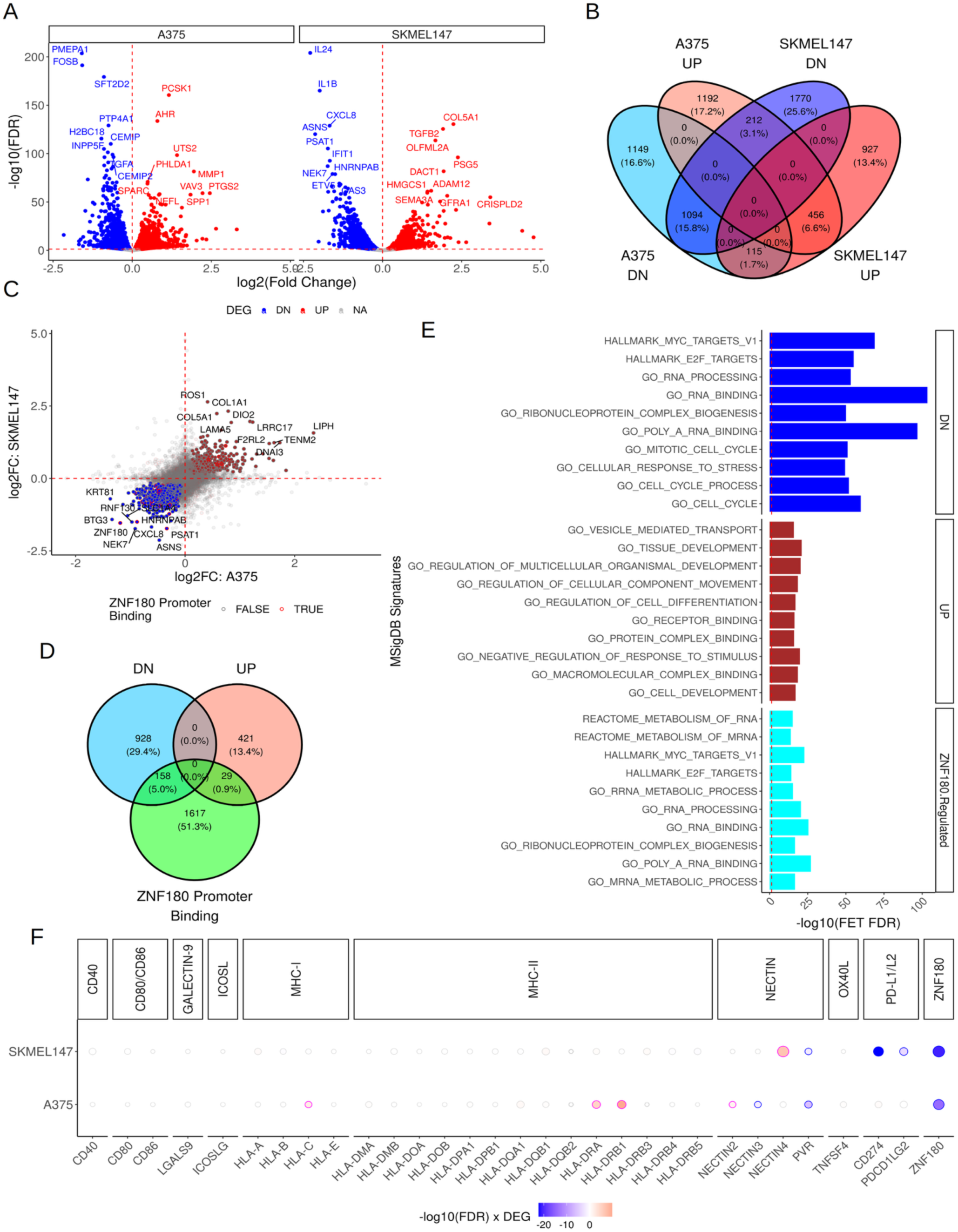
Differentially expressed genes (DEGs) and pathways by ZNF180 knockdown (KD) in A375 and SKMEL147 cells. **A. Volcano plots** of ZNF180 KD DEGs in A375 (left) and SKMEL147 (right) cells. **B. Venn diagrams of the DEGs in different cells. C. Consensus DEGs in both cell lines.** x-axis: log2(fold changes) in A375 cells, y-axis: log2(fold changes) in SKMEL147 cells. Commonly up- or down-regulated genes by FDR < 0.05 in both cells are marked in red and blue accordingly. DEGs with ZNF180 TFBS on their respective promoters are further highlighted by red borders. Top 10 up- and down-regulated genes are shown in the labels. **D. Venn diagram between the consensus *ZNF180* KD DEGs and *ZNF180* TFBS on the promoters**. The numbers of genes in different overlaps and their proportions in the union set are shown. The overlap between down-regulated (DN) and ZNF180 promoter binding genes serve as the ZNF180-regulated signature. **E**. **Enriched pathways and functions in *ZNF180* KD DEGs**. X-axis: −log10(FDR adjusted FET p-value), Y-axis: Top 10 enriched MSigDB signatures per DEG signature are shown. **F. Differentially expressed immune checkpoint genes by ZNF180 KD**. X-axis: Key immune checkpoint genes on tumor. Y-axis: A375 or SKMEL147 cells. Significant up-/down-regulated DEGs are marked by red/blue with sizes proportional to log2(fold change).

The consensus down-regulated signature was also significantly enriched for oncogenic pathways such as cell cycle, E2F targets, MYC targets and RNA processing machineries such as RNA binding proteins, spliceosome and RNA metabolic processes (**Figure 2E**; see **Supplemental Table 1B** for list of enriched pathways). Consistent with our previous study^8^, we also observed DNA repair pathways were down-regulated in the consensus signature (Fisher’s Exact Test (FET) FDR=1.55E-32, 5.73 enrichment fold change (EFC)). Interestingly, the consensus DEGs included Fanconi anemia (FA) complex genes and chromosome organization as significantly down-regulated (FA pathway: FET FDR = 3.54E-3, 7.88 EFC; chromosome organization: FET FDR = 1.15E-44, 4.63 EFC). Further, we observed significant down-regulation of a chemokine, *CXCL8* (binds to CXCR1/2), in both cell lines (Figure 2C). Tumoral CXCL8 is pro-tumorigenic, promotes migration and invasion(40–42) and is a potent chemo-attractant to immune cells including neutrophils(43), monocyte-derived suppressor cells (MDSCs)(44,45), CXCR1/2 expressing CD8 T-cells(46) and NK-cells(47).

The up-regulated genes, on the other hand, were significantly enriched for apoptosis (FET FDR = 1.14E-7, 7.22 EFC), p53 pathway (FET FDR = 4.86E-9, 7.13 EFC) and interferon-γ (IFN-γ) response (FET FDR = 4.59E-5, 5.05 EFC). The enrichment of apoptotic pathways agrees with the anti-tumor phenotypes by *ZNF180* silencing from our previous study, showing tumor regression *in vitro* and *in vivo^8^*. In addition, the up-regulation of interferon-γ response suggests increased immunogenecity in the tumor cells, for example, through promoting MHC-I and -II expressions^23,24^. Indeed, the DEGs included up-regulation HLA-C, HLA-DRA and HLA-DRB1 in A375 cells (**Figure 2F**). Interestingly, *PVR* (also known as CD155), a major ligand to the TIGIT inhibitory receptor in T-cell, was consistently down-regulated in both cell lines, and PD-L1/L2, major ligands to the T-cell inhibitory PD-1 checkpoint, were also down-regulated in SKMEL147 cells (**Figure 2F**).

Collectively, the DEGs portray pro-tumorigenic roles of *ZNF180* and its contributions to immune evasion.

### *ZNF180* silencing reshapes tumor epigenetic landscapes to reprogram into de-differentiated, immuno-suppressive melanoma through AP-1 transcription factors

We also performed ATAC-sequencing on the melanoma cells to dissect epigenetic changes by *ZNF180* silencing (see **METHODS** for **ATAC-sequencing data generation; Supplemental Figure 4**). We remark that shZ3-treated SKMEL147 cells were omitted for further analysis due to low sequencing depths (**Supplemental Figure 4D**).

*ZNF180* silencing resulted in more accessible genomic regions in both cell lines. Through identifying differentially accessible peaks (DAPs) with FDR < 0.05 (**Supplemental Table 2A, D**), we identified 4,636 peak gains and 387 peak losses in A375 cells, and 853 peak gains and 223 peak losses in SKMEL147 cells (**Figure 3A**; see **METHODS** for **ATAC-sequencing data processing**). Of these, the peak gain regions predominantly occupied promoter regions, compared to the peak loss regions (**Figure 3B**).

**Figure 3.**
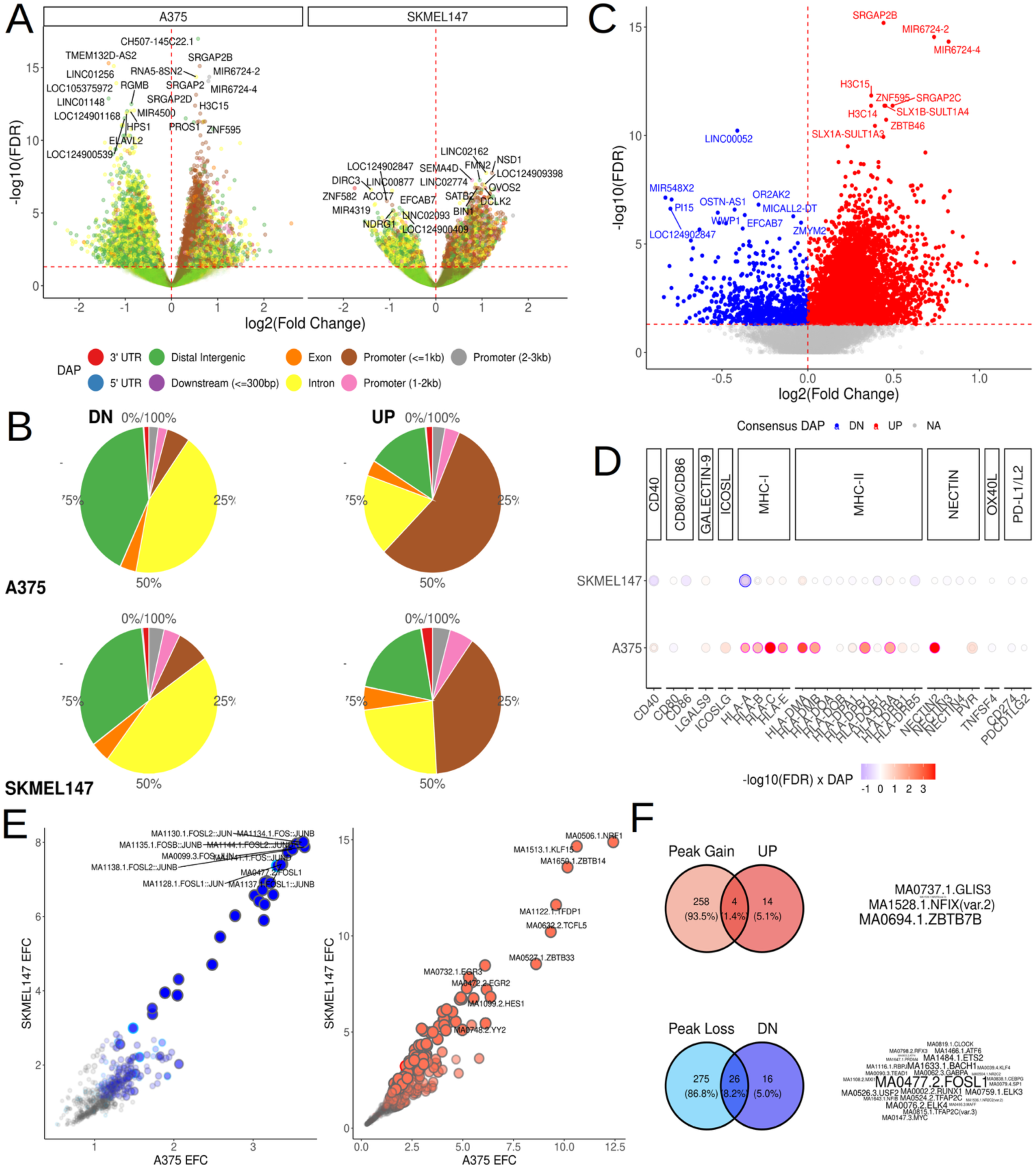
Altered epigenetic landscapes in melanoma cells by ZNF180 silencing. **A. Differentially accessible peaks (DAPs) by ZNF180 KD** in A375 (left) and SKMEL147 (right) cells. X-axis: log2(fold change) between ZNF180 silenced cells (shZ1 and shZ3) and control (NTC) cells. Y-axis: −log10(FDR). The colors denote different genomic regions as shown in the legend below. Top 10 DAPs are labeled with respective gene symbols. **B**. Proportions of different genomic regions of the DAPs in **A** (Top: A375. Bottom: SKMEL147; Left: Peak loss. Right: Peak gain). **C. Differentially regulated genes by *ZNF180* KD, shared in A375 and SKMEL147 cells.** X-axis: The average of log2(fold changes) of DAPs in promoter region of each gene across the two cell lines. Y-axis: −log10(Consensus FDR) across the two cell lines. **D**. **Differentially regulated immune checkpoint genes in melanoma.** X-axis: Key immune checkpoint genes on tumor. Y-axis: A375 or SKMEL147 cells. Significant peak gain/loss in the respective promoter regions are marked by red/blue with sizes proportional to log2(fold change). **E**. **Commonly enriched motifs in DAPs from A375 and SKMEL147 cells** (Left: Significant peak loss, Right: Significant peak gain). X-axis: Enrichment fold changes in A375. Y-axis: Enrichment fold changes in SKMEL147 cells. **F. ZNF180 regulated motifs**. Significant motifs in E were intersected with significant DEGs in Figure 2 in the Venn diagrams on the right. The motifs in the intersections are shown on the right for significant gains (top) and losses (bottom).

Through integrating the DAPs in gene promoter regions from both cell lines (**Supplemental Table 2B;** see **METHODS** for **ATAC-sequencing data processing**), we identified consensus DAPs to dissect robust functional changes in epigenetic landscapes in the melanoma cells (**Figure 3C**). Enriched pathways in the consensus peak gains reflected substantial changes in the global epigenetic landscape (**Supplemental Table 2C**) such as chromatin modifications (FET FDR = 4.67E-20, 2.32 EFC), histone binding (FET FDR = 1.84E-8, 2.21 EFC), p53 pathway (FET FDR = 6.97E-15, 2.39 EFC) and adaptive immune system (FET FDR = 2.24E-10, 1.68 EFC). In tandem, we observed significant peak gains in genes coding for MHC-I (HLA-A, -B, -C and -E) from A375 cells predominantly, and *HLA-A* in SKMEL147 cells (**Figure 3D**). Overall, these data show *ZNF180* silencing enhances the epigenetic landscape to improve the immunogenicity in melanoma cells.

On the other hand, enriched pathways in the consensus peak losses reflected suppression of neuronal pathways such as neuroactive ligand-receptor interaction (FET FDR = 9.77E-4, 2.78 EFC), synapse membrane (FET FDR = 2.64E-6, 3.58 EFC), and extracellular matrix (FET FDR = 5.65E-5, 2.64 EFC). These pathways suggest *ZNF180* silencing suppresses neural phenotypes observed in more invasive, de-differentiated neural crest-like melanoma cells(48), and *ZNF180*’s potential role to regulate melanoma plasticity.

To further understand the regulatory landscapes in the *ZNF180*-driven epigenetic changes, we systematically identified commonly enriched TF motifs in both cell lines amongst the DAPs for significant peak losses (left, **Figure 3E**) and gains (right, **Figure 3E**) (**Supplemental Table 2D**; see **METHODS** for motif enrichment analysis). The enriched motifs in the peak losses captured Activator Protein 1 (AP-1) family of TFs such as *FOS, FOSL1/2* and *JUN/JUNB*(49,50). Of these, several enriched TFs such as *FOSL1* and MYC were also significantly down-regulated by *ZNF180* silencing (lower, **Figure 3F**), indicating that *ZNF180* is an upstream regulator of their TF activities in melanoma cells.

In melanoma, the AP-1 network drives reprogramming of melanoma cells and determines the melanoma differentiation states(49). Particularly, up-regulation of *FOSL1*, as the most enriched motif among the significantly down-regulated TFs by *ZNF180* silencing (**Figure 3E, F**), is oncogenic and reprograms melanocytes through down-regulation of *MITF* in an HMGA1-dependent manner(51). The differential expressions by *ZNF180* silencing supports this through showing significant *MITF* up-regulations (A375 cells: FDR=4.00E-2, 1.13-fold increase), and HMGA1 down-regulations (SKMEL147 cells: FDR=9.47E-50, 0.49-fold decrease, A375 cells: FDR=3.88E-29, 0.77-fold decrease) (**Figure 3F**).

Another example of AP-1 network mediated melanoma reprogramming involves c-Jun (also known as JUN). The reciprocal antagonism between *MITF* and c-Jun through competitive binding for c-Jun regulated genomic regions. *MITF* reduction increases c-Jun and TNF-stimulated cytokine expressions to yield de-differentiated, pro-inflammatory melanoma with increased immuno-suppressive myeloid recruiments(52). Indeed, *ZNF180* KD significantly down-regulated the TNFα signaling pathway in melanoma cells *in vitro* (TNFA SIGNALING VIA NFKB: FET FDR = 2.30-E-6, 3.68 EFC; **Supplementary Table 1B**).

### *ZNF180*-regulome underlies AXL-biased, de-differentiated melanoma fate determination

Beyond AP-1 TFs, we identified *ZNF180*-regulated protein-protein interaction (PPI) network landscape to dissect the functional influences of the consensus DEGs and DAPs (**Figure 4A**). Utilizing the high confidence PPIs (confidence score > 0.9) from STRING database(53), we gathered the overlapping proteins in the database with the consensus DEGs and DAPs (**Figure 4B**) to extract the connected PPI network as *ZNF180*-regulome (see **Methods** for **STRING PPI analysis of ZNF180-regulome; Supplemental Table 3**).

**Figure 4.**
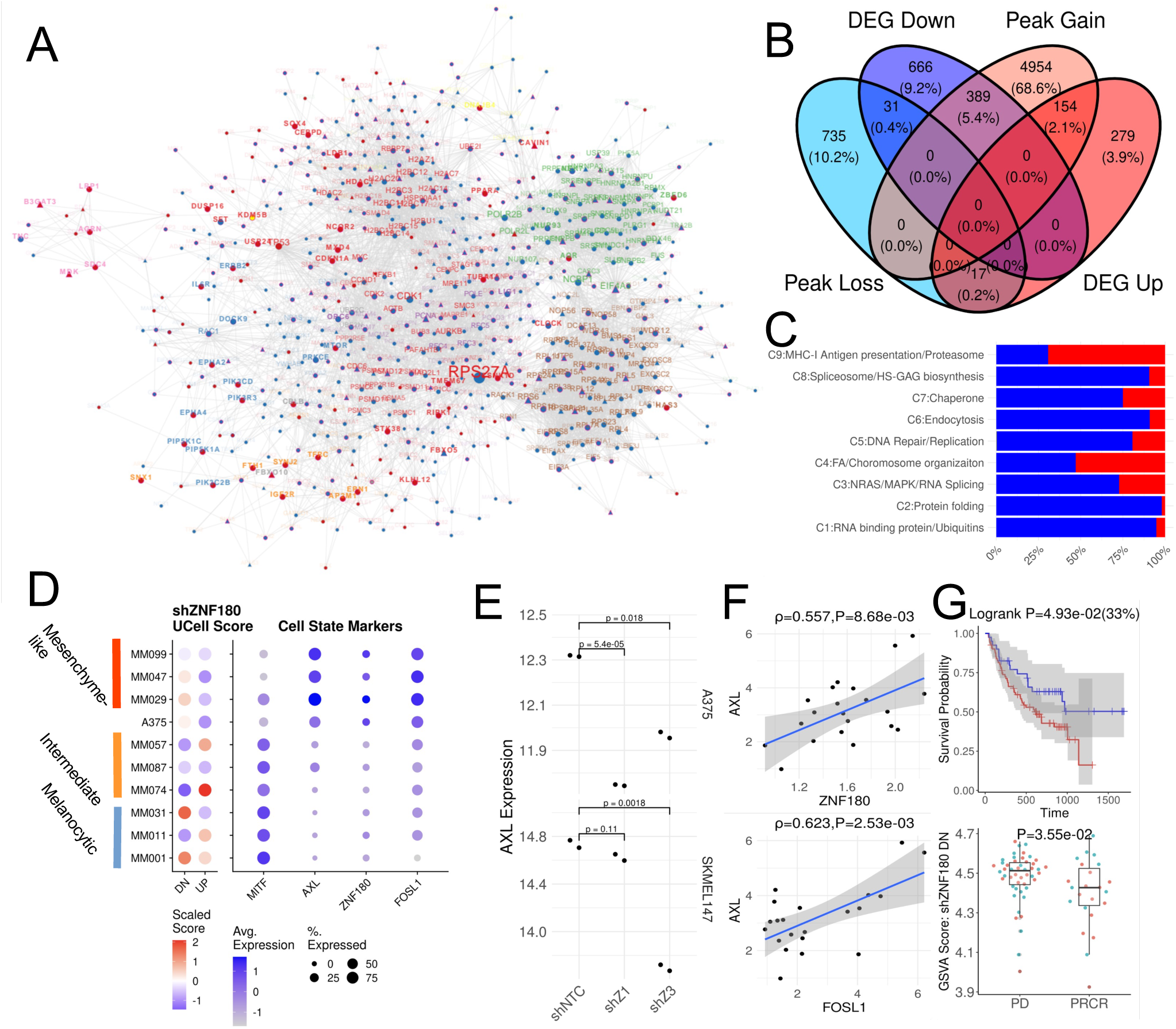
*ZNF180*-regulome drives tumors towards invasive, therapy-resistant low MITF/AXL ratio states and predicts immunotherapy response. **A. Dys-regulated protein-protein interaction (PPI) network by *ZNF180* silencing.** Significant up-/down-regulated DEGs were colored red/blue in the nodes, and significant gain/loss DAPs were colored magenta/purple on the node borders. Triangle nodes are genes whose promoters overlap with ZNF180 TFBS. Gene labels are color-coded for different subnetworks as shown in the legend. Bold labels highlight down-/up-regulated DEGs whose promoters overlapped with significant gain/loss DAPs in accessibility. B. **Venn diagram of signifcant DEGs and DAPs**. **C**. Proportions of up- and down-regulated DEGs in each subnetwork in red and blue respectively. **D. ZNF180-regulated pathways in single-cell transcriptomes of different melanoma cells**. <u>Left:</u> The dotplot shows overall activations/suppressions of up- (UP) or down-regulated (DN) signatures by *ZNF180* silencing in different melanoma cells. <u>Right</u>: The dotplot shows overall expressions of *MITF*, *AXL*, *ZNF180* and *FOSL1* in single-cell transcriptomes of melanoma cells. The melanoma cells are stratified by the subtype classifications in Wouters *et al.* 2020. **E, F. AXL regulation by *ZNF180-FOSL1* axis**. *AXL* down-regulation by *ZNF180* silencing in A375 (top) and SKMEL147 (bottom) is shown in **E**. *AXL* is significantly correlated to *ZNF180* (top) and *FOSL1* (bottom) in ipilimumab-treated metastatic melanoma from Snyder *et al.* 2014 cohort as shown in **F**. **G**. ***ZNF180*-regulome signature is predictive of anti-PD-1 responses**. <u>Top</u>: Kaplan-Meier plot to show poor prognosis in patients with high ZNF180 regulation score by median from Liu *et al.* 2019 cohort. <u>Bottom</u>: Boxplot to show significantly higher ZNF180 regulation scores in the non-responsive progressive disease (PD) group, compared to the responsive partial response or complete remission (PR/CR) group from Riaz *et al.* 2017 cohort.

Overall, we identified 9 coherently connected modules of *ZNF180*-regulome with compartmentalized functions and pathways (**Figure 4C; Supplemental Figure 5**). We observed up-regulation of Fanconi Anemia (FA)-complex/chromosome organization in C4, which aligns with close interactions between *ZNF180* and FA complex and dys-regulations in DNA repair pathways from our previous study on the gene network model of primary melanoma(8). Further, we observed up-regulation of MHC-I antigen presentation in C9. This coincides with one of aggressive features of de-differentiated AXL-high tumor as immune-cold tumors through MHC-I suppression while undergoing Epithelial-Mesenchymal transition (EMT)(39,54).

On the other hand, RNA-bidning ubiquitins (C1), protein folding (C2), endocytosis (C6), spliceosome/HS-GAG biosynthesis were among the most down-regulated modules. Again, these pathways closely align with our previous findings on the primary melanoma network model(8), and include several pathways driving EMT. One example is aberrant spliceosome, known to induce EMT-inducing transcription factors such as *ZEB1*(55) whose expressions were robustly down-regulated by *ZNF180* silencing (A375: FDR=1.20E-2, 0.897-fold decrease; SKMEL147: FDR=8.07E-17, 0.622-fold decrease).

In tandem, we observed that *ZNF180* orchestrates the master regulators of melanoma lineage differentiation. *ZNF180* was over-expressed in mesenchyme-like melanoma cells, concurrently with down-regulation of *MITF*, and up-regulations of *AXL* and *FOSL1* from the single-cell transcriptomes of patient-derived melanoma cell lines(56) (**Figure 4D**). Particularly, we robustly observed *ZNF180* silencing robustly down-regulates *AXL* in A375 and SKMEL 147 cells, indicating ZNF180 is an upstream regulator of *AXL*, and may drive AXL-biased melanoma lineages (**Figure 4D, E**). Indeed, we observed significant correlations between *AXL* and *ZNF180,* and *AXL*, and *FOSL1* (*ZNF180*-driven TF) and *AXL* in bulk transcriptome of untreated metastatic melanoma from Snyder *et al.* 2014(11) (**Figure 4F**). These altogether indicate *ZNF180* drives melanoma plasticity in primary tumors through *FOSL1*-modulated cellular reprogramming, leading to MHC-I low, MITF^low^/AXL^high^ de-differentiated phenotype in anti-PD-1 resistant melanoma(39).

### *ZNF180* silencing alters immune-suppressive tumor microenvironment

*ZNF180* silencing altered T-cell immune checkpoint ligand profiles in the tumors. We performed Western blotting for various T-cell immune checkpoint ligands on A375 and SKMEL147 cells with or without IFN-γ stimulation *in vitro* (**Supplemental Figure 6**; see **METHODS** for **Western blotting**). Among these ligands, CD155 (also known as PVR), the primary ligand for TIGIT inhibitory receptor on T-cells to modulate PD-1/PD-L1 ICI resistance(57), was most robustly suppressed by *ZNF180* silencing in both cells and regardless of IFN-γ stimulation (**Figure 5A**). In SKMEL147 cells, we also observed suppression of PD-L1 and Galectin-9 regardless of IFN-γ (**Supplemental Figure 6A**). In A375 cells, PD-L1 expressions were present only in IFN-γ stimulation, and this was also suppressed by *ZNF180* silencing (**Supplemental Figure 6B**).

**Figure 5.**
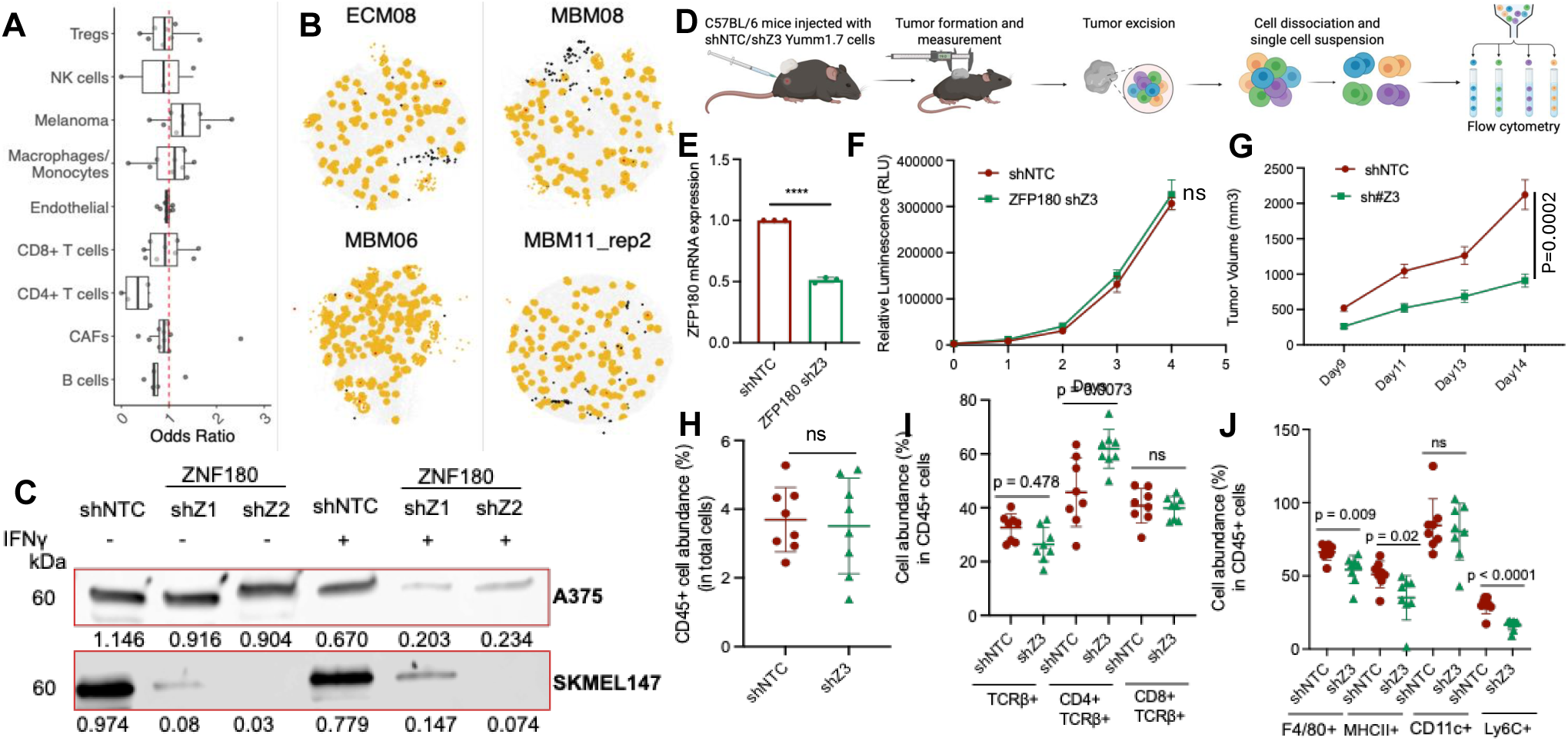
*ZNF180* as driver of CD4 T-cell mediated cytotoxicity. **A. ZNF180+ tumor neighborhood in spatial transcriptomes**: Enrichments (odds ratio > 1) or depletion (odds ratio < 1) of various cell types in ZNF180+ tumor neighborhoods from Biermann *et al.* 2022 (Red: extracranial metastasis (ECM), green: bone metastasis (MBM)). **B. Exemplary Spatial transcriptomes** showing depleted CD4 T-cells (black) in ZNF180+ tumor (red) and its neighborhoods (yellow). **C. Western blots (WB) of CD155 shows CD155 suppression by ZNF180 silencing**. Different shRNAs (shNTC: non-transfected control, shZ1, shZ2: shZNF180) are shown on the top. WB staining results without and with IFN-γ stimulation are shown on left and right respectively for A375 (top) and SKMEL147 (bottom) cells. **D-J. *ZNF180* silencing alters immune cell landscapes in immune competent murine model. D.** Schematic of the in vivo experimental workflow. **E**. Quantitative RT-PCR analysis of ZNF180 mRNA expression in Yumm1.7 cells transduced with non-targeting control shRNA (shNTC) or ZNF180-targeting shRNA (shZ3). Expression is normalized to GAPDH. Data are shown as mean ± SEM. ****p < 0.0001, unpaired t-test. **F**. *In vitro* proliferation of shNTC and shZFP180 Yumm1.7 cells measured by luminescence-based viability assay over 4 days. Data represent mean ± SEM; ns, not significant. **G**. Tumor growth kinetics of Yumm1.7 cells (shNTC vs. shZFP180) subcutaneously implanted in C57BL/6 mice. Tumor volumes were measured every 3–4 days. Data represent mean ± SEM. **H**. Total CD45⁺ immune cell abundance as a percentage of all live cells in tumors from shNTC or shZ3 Yumm1.7-bearing mice. No significant difference observed. **I**. T cell subset analysis showing percentages of total TCRβ⁺, CD4⁺TCRβ⁺, and CD8⁺TCRβ⁺ populations among CD45⁺ cells. shZ3 tumors exhibited a significant increase in CD4⁺TCRβ⁺ cells (p = 0.0073) but no significant differences in total TCRβ⁺ or CD8⁺TCRβ⁺ subsets. **J**. Myeloid cell composition among CD45⁺ cells, including F4/80⁺ macrophages, MHC II⁺ antigen-presenting cells, CD11c⁺ dendritic cells, and Ly6C⁺ monocytes. *Zfp180* silencing significantly reduced Ly6C⁺ cells (p < 0.0001), MHCII⁺ (p = 0.009), and CD11c⁺ (p = 0.02) populations, with no significant difference in F4/80⁺ macrophages. Data represent individual mice; bars indicate mean ± SEM. Statistical comparisons were performed using unpaired two-tailed t-tests; *p < 0.05, **p < 0.01, ****p < 0.0001, ns = not significant.

In tandem, *ZNF180*+ tumor neighborhoods in metastatic melanoma showed CD4 T-cell exclusions. We systematically analyzed spatial transcriptomes of extracranial and bone metastatic melanoma (ECM & MBM) to dissect immune cell landscapes in ZNF180+ tumor neighborhoods at near cellular resolution from Biermann *et al.* 2022(58) (see **Methods** for **Biermann *et al.* 2022 spatial transcriptome analysis**). Most notably, *ZNF180*+ tumor neighborhoods exhibited robust depletion of CD4 T-cells (i.e. odds ratio < 1) and moderate enrichments of macrophages (i.e. odds ratio > 1) across multiple samples of metastatic melanoma spatial transcriptome from Biermann *et al.* 2022(58) (**Figure 5A, B**; see **Methods** for **Biermann *et al.* 2022 spatial transcriptome analysis**).

We validated *ZNF180* silencing alters immune cell landscape and modulates tumor killing by tumor extrinsic factors *in vivo*. These were validated by *Zfp180* knock-down (murine ortholog to human *ZNF180*) in immune competent Yumm 1.7 xenograft to syngenic C57BL/6J murine model (**Figure 5D**; See **Methods** for ***In vivo* xenografts**). Knockdown of *Zfp180* was validated by qRT-PCR, confirming a robust reduction in mRNA expression compared to control (****p < 0.0001) (**Figure 5E**). *In vitro* proliferation assays showed no significant difference in growth between *Zfp180*-silenced control Yumm1.7 cells (**Figure 5F**). Further, tumor growth analysis revealed no significant differences in early tumor growth; however, tumor volume measurements showed significantly reduced progression in *Zfp180*-silenced tumors by day 14 (p = 2E-4) *in vivo* (**Figure 5G-H**). Flow cytometry demonstrated that *Zfp180* silencing selectively reshaped the tumor microenvironment, with no changes observed in splenic immune cell populations (**Supplemental Figure 7-9**), confirming the effect was tumor-localized. Within tumors, overall CD4⁺ and CD8⁺ T-cell frequencies were unchanged, but a significant increase in TCRβ⁺ CD4⁺ T cells was detected (p = 7.3E-3), while TCRβ⁺ CD8⁺ T cells remained unaffected (**Figure 5I, Supplementa**l **Figure 10**). Further, we observed significant decrease in NK cells (p = 3E-4;**Supplementa**l **Figure 11**). On the myeloid side, macrophages (F4/80⁺) were significantly decreased (p = 9E-3), accompanied by reduced MHCII⁺ antigen-presenting cells (p = 2E-2) and a striking loss of Ly6C⁺ inflammatory monocytes (p < 1E-4). CD11c⁺ dendritic cells, however, were not significantly affected (**Figure 6K**). Together, our results demonstrate that Zfp180 loss enhances CD4⁺ helper T-cell recruitment while suppressing multiple myeloid populations, including macrophages and Ly6C⁺ cells, thereby shifting the tumor immune landscape toward a more T-cell–dominant environment.

**Figure 6.**
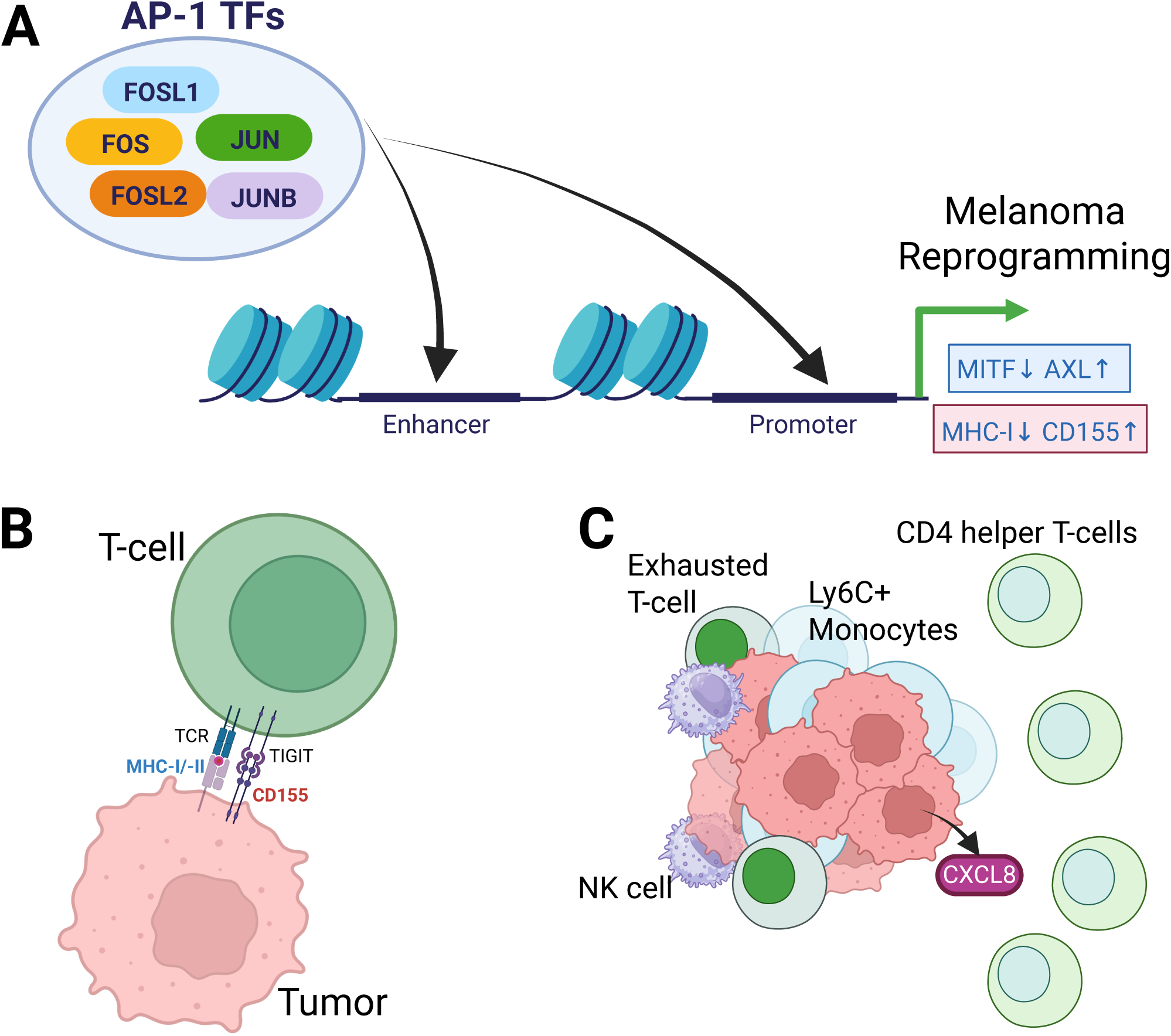
Overview of ZNF-regulome in melanoma and its immune-suppressive features. **A. *ZNF180* promotes melanoma reprogramming through enhancing AP-1 TF activities**. *ZNF180* altered the melanoma genome accessibility landscapes to promote AP-1 TF activities and modulate melanoma reprogramming (higher AXL/MITF ratio), loss of immunogenicity (MHC-I loss) and T-cell inhibitory signals (CD155 gain). **B**. ***ZNF180* evades T-cell immune surveillance** through MHC-I loss (colored blue), and relaying T-cell inhibitory signals through TIGIT/CD155 checkpoint (colored red). **C**. ***ZNF180* fosters immune-suppressive tumor microenvironment.** Upon ZNF180 silencing, tumors microenvironment undergoes changes in immune microenvironments (highlighted by blue arrow) to exclude pro-inflammatory Ly6C+ monocytes and NK-cells and recruit CD4 helper T-cells. These changes could be facilitated through subsequent *CXCL8* silencing (highlighted by red arrow), a pro-tumorigenic chemokine regulated by *ZNF180* within tumor cells.

## DISCUSSION

### *ZNF180* modulates tumor intrinsic proto-oncogenic pathways

Our findings from the previous study were robustly recapitulated through confirming consistent regulatory roles of *ZNF180* on several pro-tumorigenic pathways. Upon *ZNF180* silencing via shRNA knock-down on melanoma cells with different genetic backgrounds (A375: BRAF-mutant, SKMEL147: NRAS-mutant), we observed consistent down-regulation of protooncogenic pathways such as MYC targets(59), RNA splicing(60) and DNA repair pathways during replication such as Fanconi-Anemia (FA) complex and MSH2(61,62) (**Supplemental Table 1A, B; Figure 2E**), and they constituted closely interacting PPI subnetworks (**Figure 4A, C**). These pathways are known to be associated with relapse and metastasis(63–66), and collectively, activation of these *ZNF180* signatures were robustly associated with conferring ICI resistances (**Figure 1D**, **Figure 4G**).

These *ZNF180*-regulated pathways included potential therapeutic avenues to abolish tumor intrinsic ICI resistance factors. While MYC and RNA splicing have been implicated as established drivers of ICI resistance(59,64,67), the roles of FA complex in ICI resistance are yet unknown. FA complex‘s primary function is to recognize and repair DNA interstrand crosslinks (ICLs) during S phase in cell cycle and allow accurate DNA synthesis using the homologous chromosome as a template (**Figure 6A**). Its dysregulation promotes mitosis with unrepaired DNA damages and introduce genomic instability(61,62). While defective FA complex is often associated with better responses to immunotherapy with increased immunogenecity and neoantigens(68), its up-regulations in tumor cells have been linked to maintain DNA integrity in these tumor cells and promote drug resistance(69). Herein, *ZNF180*-modulated FA complex up-regulation renders a novel potential mechanism to confer ICI resistance in early stage melanoma and FA core complex as a new therapeutic avenue, for instance, by targeting the FANC-D2-I heterodimer as a unique substrate(70).

### *ZNF180* regulates AP-1 transcriptions factors to drive melanoma plasticity and emergence of immune-suppressive de-differentiated tumors

*ZNF180* showed direct regulations towards AP-1 transcription factors (TFs) underlying melanoma plasticity and reprogramming to mesenchyme-like melanoma(50,51,71) (**Figure 6B**). AP-1 is the major regulator of mesenchyme-like state enhancers in melanoma(72,73), and *ZNF180* showed epigenetic regulations on the TFs. *ZNF180* knock down significantly reduced accessible genomic regions enriched for DNA binding motifs for AP-1 TFs including *FOS, FOSL1/2, JUN* and *JUNB* (**Figure 3E**).

Of those, we identified *FOSL1* as the most prominently regulated TF by *ZNF180* (**Figure 3F**). *FOSL1* is oncogenic and reprograms melanocytes through down-regulation of *MITF* in HMGA1-dependent manner(51) for which we confirmed the up-regulations of *MITF* and *HMGA1* by *ZNF180* silencing. We further observed regulatory roles of *ZNF180* in driving the melanoma reprogramming through regulating *AXL* expression (**Figure 4E, F**). These concomitantly indicate *ZNF180* drives melanoma reprogramming towards AXL^high^/MITF^low^ state, a defining feature of de-differentiated melanoma.

De-differentiated melanoma is an invasive melanoma subtype(39,74) with immune-suppressive features including loss of MHC-I(75), up-regulation of inhibitory checkpoint ligands such as PD-L1(76) and facilitate T-cell exclusion or dysfunction(76). As discussed in the later sections, these immune-suppressive features such as MHC-I loss and inhibitory T-cell signaling were reproduced by *ZNF180* in this study. Taken together, *ZNF180* is a novel master regulator of melanoma plasticity, leading to immune-suppressive phenotypes in these tumors.

### *ZNF180* modulates key changes in immune-modulatory signals in genetic-dependent manner

*ZNF180* silencing improved immunogenicity melanoma cells, primarily in *BRAF*-mutant A375 cells. *ZNF180* silencing rescued MHC-I expressions predominantly (**Figure 2F**, **Figure 3D**), but it remained unchanged in *NRAS*-mutant SKMEL147 cells. The observed MHC-I changes in A375 cells were in alignments to the MHC-I loss in anti-PD-1 resistant *ZNF180*+ tumors in Jerby-Arnon *et al.* 2018 scRNA-seq (**Figure 1F-H**), and up-regulated MHC-I surface expressions upon *ZNF180* knock-out in IFN-γ stimulated B16 cells (**Figure 1I,J**).

Also, *ZNF180* impacted several key ligands for T-cell inhibitory checkpoints, and some of the changes were unique to each cell line. Ligands to PD-1, PD-L1 and -L2, were significantly down-regulated in *NRAS*-mutant SKMEL147 cells by *ZNF180* knock down while PD-L1 and PD-L2 expressions had little changes in *BRAF*-mutant A375 cells (**Figure 2F**, **Figure 3D**). Indeed, *NRAS*-mutant melanoma are known to have higher PD-L1 expressions, hence show better responses to PD-1 blockades in advanced melanoma(77).

These results suggest that immune-suppressive functions of *ZNF180* is dependent on the genetic backgrounds: its impacts on *BRAF*-mutant tumors favors loss of immunogenicity to cloak the tumors from anti-tumor immune cells, while *ZNF180* could favor resistance to lysis through modulating T-cell inhibitory signals in *NRAS*-mutant cells. These differences warrant future studies to investigate robustness of the findings in more cells with known genetic backgrounds.

### *ZNF180* fosters immune-suppressive microenvironment through promoting TIGIT/CD155 T-/NK-cell inhibitory signals and helper T-cell exclusion

We report that *ZNF180* is an upstream regulator of tumoral CD155 through their robust co-expressions in anti-PD-1 resistant tumors (**Figure 1H**), patient-derived melanoma cell lines (**Figure 4D**), and robust down-regulation by ZNF180 KD *in vitro* (**Figure 2F**, **Figure 5A, B**). CD155 is the primary ligand to the inhibitory TIGIT receptor to trigger inhibitory signals in T- and NK-cells(78,79). Particularly, TIGIT/CD155 checkpoint mediates ICI resistance in melanoma(57,80) through inhibiting anti-tumor T-cell effector functions(81) and inducing immunosuppressive DCs(82). CD155 also binds to the co-stimulatory receptor CD226 (also known as DNAM-1), yielding competitive binding for CD155 between these two receptors to regulate T-cell activation/inhibition balance(83), anti-tumor CD8 T-cell effector functions(81) and pro-inflammatory (Th1/Th17)/anti-inflammatory(Th2) balance(84).

In the light of the intricate balance between TIGIT and CD226 for CD155 binding, *ZNF180* favored T-cell inhibitory signals through CD155 in TIGIT dominated immune landscape. Using Liu *et al.* 2019 bulk transcriptomes from responders and non-responders of anti-PD-1 therapy(36), *ZNF180* expressions were significantly correlated with the inhibitory TIGIT/CD155 checkpoint expressions while the correlations with the co-stimulatory CD226/CD155 checkpoints were not significant amongst the non-responders (**Supplemental Figure 12**).

Further, *ZNF180* regulated TCRβ+ CD4 T-cells infiltrations in tumor microenvironment. While traditionally known as “helper” cells that orchestrate the immune response, TCRβ+ CD4 T-cells can foster anti-tumor immune microenvironment. Helper CD4 T-cells licenses dendritic cells to assist activation of naïve CD8 T-cells(85), releases essential cytokines to modulate the effector actions of cytotoxic CD8 T-cells and NK-cells(86), and can differentiate into cytotoxic T lymphocytes that directly kill cancer cells(87–89).

These results suggest *ZNF180* collectively modulates immune-suppressive tumor microenvironment by activating the T-cell inhibitory checkpoints (**Figure 6C**) and excluding the helper T-cells (**Figure 6D**), thereby lack of their functions to promote anti-tumor immune microenvironment.

### *ZNF180* silencing modulates recession in NK-cells and myeloid cell subsets potentially through *CXCL8*

We remark that *ZNF180* silencing significantly reduced NK-cells (**Supplemental Figure 11**), macrophages (F4/80+) and inflammatory monocytes (Ly6C+) in the tumors (**Figure 5J**) in immune competent Yumm 1.7 xenograft mouse model. To this end, down-regulation of *CXCL8* (also known as IL-8) by *ZNF180* silencing renders potential explanations to the associated changes in the immune microenvironment. As the ligand that binds to CXCR1/2, *CXCL8* has been shown to be expressed in lymph node metastatic melanoma and facilitated recruitment of highly cytotoxic CD56dimKIR+CD57+ NK cells(90). Also, *CXCL8* is a potent chemo-attractant to myeloid-derived suppressor cells to produce immune-suppressive, inflammatory tumor microenvironment including modulation of JAK/STAT3 activation(91), inhibition of anti-tumor T-cell responses, promotion of tumor metastasis and growth(41,42). The co-expression of *ZNF180* and *CXCL8-JAK/STAT3* was also observed in mesenchymal-like melanoma cells predominantly (**Supplemental Figure 13**). This evidence suggests *CXCL8* suppression by *ZNF180* silencing could facilitate recession of immune-suppressive myeloid subsets to foster anti-tumor T-cell activities in the expense of reducing NK-cell anti-tumor activities. Further, the exclusion of NK-cells through *CXCL8* suppression could limit the effects of CD155/TIGIT checkpoint inhibition within the T-cells, hence selectively enhancing T-cell dominated microenvironments. Taken together, CXCL8-CXCR1/2 axis provides a potentially promising interventional therapeutic avenues(40) for *ZNF180* modulated immunotherapy resistance and these warrant further investigations in the future studies.

## CONCLUSION

*ZNF180* is a multi-facetted tumor intrinsic regulator to foster immune-suppressive tumor microenvironment at multiple levels. Within tumors, *ZNF180* reprograms melanoma cells to exhibit invasive, immune-cold phenotypes with helper T-cell exclusions to evade immune surveillance. Our study proposes potential therapeutic strategies towards *ZNF180*-driven tumors to intervene their ICI resistance through targeting its downstream factors such as FA core complex and TIGIT/CD155 blockade, and *ZNF180*-regulome silencing as a novel interventional strategy for non-metastatic melanoma.

## LIST OF ABBREVIATIONS

CPM: Counts Per Million
CR: Complete remission
EDTA: Ethylenediaminetetraacetic acid
DAP: Differentially accessible peaks
DEG: Differentially expressed genes
DN: Down-regulation
DOI: Digital Object Identifier
EFC: Enrichment fold change
EMT: Epithelial-mesenchymal transition
ENCODE: the Encyclopedia of DNA Elements
FA: Fanconi-Anemia complex
FACS: Fluorescence-activated cell sorting
FBS: Fetal bovine serum
FDR: False discovery rate
FET: Fisher’s Exact Test
GEO: Gene Expression Omnibus
ICI: Immune checkpoint inhibition
KD: knock-down
kNN: k-nearest neighbor
KO: knock-out
MACS: Model-based Analysis of ChIP-seq
IFN-g: Interferon-gamma
OR: Odds ratio
PBS: Phosphate-buffered saline
PC: Principal component
PD: Progressive disease
PE: Paired-end
PPI: Protein-protein interaction
PR: Partial response
QC: Quality control
RPKM: Reads per Kilobase Million
SD: Stable disease
ssGSEA: Single-sample Gene Set Enrichment Analysis
TF: Transcription factor
TFBS: Transcription factor binding site
TBST: Tris-buffered saline with 0.05% Tween-20
TCGA: The Cancer Genome Atlas
TME: Tumor microenvironment
TMM: Trimmed mean of M-values
TSS: Transcription starting site
UP: Up-regulation

## DECLARATIONS

### Ethical Approval

Not applicable.

### Competing Interests

The authors have no relevant financial or non-financial interests to disclose.

### Author contributions

Conceptualization: W.M.S. and P.A.; Methodology: W.M.S. and P.A.; Data Curation: W.M.S.; Visualization: W.M.S., P.A., S.K.K., S.H.C.; Writing, Original Draft: W.M.S., X.Z., P.A., S.K.K., S.H.C.; Writing, Review & Editing: W.M.S., X.Z., P.A., S.K.K., S.H.C; Investigation: W.M.S., X.Z., P.A., Supervision: W.M.S., P.A.; Funding acquisition: W.M.S., P.A., S.H.C.

### Funding

This study was supported by the National Institutes of Health (NIH) under award numbers R35GM142918.

### Availability of data and materials

The sources of publicly available data utilized in this study are as follows:

- <u>Jerby Arnon *et al.* 2018 single-cell transcriptome:</u> We downloaded the raw count matrix and cell/sample meta data from Gene Expression Omnibus (GEO) with accession ID, GSE115978.
- <u>Biermann *et al.* 2022 spatially resolved transcriptome:</u> We downloaded 16 samples of spatial transcriptome data by SlideSeq-V2 from GSE185386
- <u>Snyder *et al.* 2014 bulk transcriptome:</u> We accessed log2-transformed, Reads per Kilobase Million (RPKM) normalized data across 21 samples from cBioPortal (https://www.cbioportal.org/study/summary?id=skcm_mskcc_2014).
- <u>Riaz *et al.* 2018 bulk transcriptome:</u> Raw gene count matrices were obtained from the published study through Gene Expression Omnibus data portal (GEO accession: GSEGSE91061).
- <u>Liu *et al.* 2019 bulk transcriptome:</u> Transcripts Per Million (TPM)-normalized values from the published study in Supplementary Data 2 were downloaded.

The sequencing data generated from this study have been made available:

- <u>ATAC- and RNA-sequencing data from *ZNF180* knockdown in A375 and SKMEL147:</u> The sequencing data have been shared through Gene Expression Omnibus (GEO) under the accession numbers (RNA-seq: GSE307748, ATAC-seq: GSE307749).

Codes to reproduce the results have been released through Zenodo with Digital Object Identifier (DOI: https://doi.org/10.5281/zenodo.17050434). The companion set of processed data have been released through Synapse (synID: syn68871742, DOI: https://doi.org/10.7303/syn68871742).

### Competing interests

The authors declare that they have no competing interests.

## Acknowledgements

This work was supported in part through the computational and data resources and staff expertise provided by Scientific Computing and Data at the Icahn School of Medicine at Mount Sinai and supported by the Clinical and Translational Science Awards (CTSA) grant UL1TR004419 from the National Center for Advancing Translational Sciences. Research reported in this publication was also supported by the Office of Research Infrastructure of the National Institutes of Health under award number S10OD026880 and S10OD030463. The content is solely the responsibility of the authors and does not necessarily represent the official views of the National Institutes of Health.

## SUPPLEMENTAL FIGURE

**Supplemental Figure 1.** Raw fluorescence intensity measurements of MHC-I in B16-F12 cells in sgRNA transfected cells (red), control gRNA transfected cells (blue) and isotype control (black).

**Supplemental Figure 2. ZNF180 suppressions in A375 (A) and SKMEL147 (B) cells by two shRNA constructs (shZ1, shZ2).**

**Supplemental Figure 3. Gene expressions by RNA-sequencing. A, B.** log2(CPM) expressions on different samples from A375 (**A**) and SKMEL147 (**B**) cells. **C, D**. Barplots to show library read depths on different samples from A375 (**C**) and SKMEL147 (**D**) cells.

**Supplemental Figure 4. A, B. Boxplots of samplewise raw read counts from ATAC-sequencing:** A375 (A) and SKMEL147 (B) cells**. C, D. Barplots of samplewise read depths from ATAC-sequencing:** A375 (C) and SKMEL147 (D).

**Supplemental Figure 5. Top 5 most enriched MsigDB pathways and functions in ZNF180-regulome modules.**

**Supplemental Figure 6. Protein expression of various ligands for T-cell immune checkpoints in ZNF180-silenced (A) SKMEL147 and (B) A375 human melanoma cells.** *ZNF180* silencing resulted in robust reduction of CD155 in both cells, as shown by Western Blot (WB). Cells with IFN-γ+ were treated with 5 ng/ml IFN-γ for 36 hours. GAPDH was used as a loading control as shown on the bottom.

**Supplemental Figure 7. Flow cytometric analysis of tumor-infiltrating lymphocytes (TILs).** Representative gating strategy showing immune subsets isolated from tumor tissues. Live single cells were first gated for CD45⁺ cells, followed by analysis of CD4⁺ T cells, CD8⁺ T cells, TCRβ⁺ T cells, and NK1.1⁺ natural killer cells. Within the TCRβ⁺ population, proportions of TCRβ⁺CD4⁺ helper T cells and TCRβ⁺CD8⁺ cytotoxic T cells were quantified. Percentages shown in each gate indicate the fraction of positive cells within the parent population.

**Supplemental Figure 8. ZNF180 silencing does not alter systemic lymphoid composition in the spleen. A.** Quantification of immune cell subsets in spleens from tumor-bearing mice injected with Yumm1.7 shNTC or shZ3 cells. The frequency of CD4⁺, CD8⁺, NK1.1⁺, TCRβ⁺, CD4⁺TCRβ⁺, and CD8⁺TCRβ⁺ populations among splenic CD45⁺ cells was measured by flow cytometry. No statistically significant differences were observed between groups. Data are shown as mean ± SEM. **B, C.** Representative flow cytometry plots showing gating strategies for major immune subsets (**B**) and TCRβ⁺ CD4+ and CD8+ T-cells (**C**) in the spleens of mice bearing shNTC (top row) and shZ3 (bottom row) tumors.

**Supplemental Figure 9. ZNF180 silencing does not alter systemic myeloid composition in the spleen. A.** Quantification of myeloid immune cell subsets among CD45⁺ splenocytes from mice bearing shNTC or shZ3 Yumm1.7 tumors. Frequencies of F4/80⁺ macrophages, CD11c⁺ dendritic cells, MHCII⁺ antigen-presenting cells, and Ly6C⁺ monocytes were assessed by flow cytometry. No statistically significant differences were observed between groups. Data are shown as mean ± SEM. **B**. Representative flow cytometry dot plots illustrating gating for each population in spleens from shNTC (top row) and shZ3 (bottom row) tumor-bearing mice.

**Supplemental Figure 10. A.** Representative flow cytometry gating strategy and dot plots for immune profiling in Yumm1.7 tumors for lymphoids (**A**) and myeloids (**B**).

**Supplemental Figure 11.** Flow cytometric quantification of CD4⁺ T cells, CD8⁺ T cells, and NK cells (CD3⁻NK1.1⁺) among CD45⁺ cells. *Zfp180* knockdown significantly reduced NK cell abundance (p = 0.0003) with no significant change in CD4⁺ or CD8⁺ T cells.

**Supplemental Figure 12. Correlations between *ZNF180* and TIGIT/CD155, DNAM-1/CD155 checkpoints. A. C**orrelations with *PVR* (i.e. CD155), **B.** Correlations with *CD226* (i.e. DNAM-1), and **C**. Correlations with TIGIT in C for non-responders (PD) and responders (SD, CR) from Liu *et al.* 2019 study(36).

## SUPPLEMENTAL TABLE

**Supplemental Table 1. Differentially expressed genes by ZNF180 knock down. A.** List of differentially expressed genes (DEGs) by *ZNF180* knock-down in A375 and SKMEL147 cells, and **B.** Enriched pathways and functions in the consensus DEGs across A375 and SKMEL147 cells.

**Supplemental Table 2. Differentially accessible peaks by *ZNF180* knock down. A.** List of differentially accessible peaks (DAPs) by ZNF180 knock-down in A375 and SKMEL147 cells, B. List of consensus DAPs from A375 and SKMEL147 cells, **C.** Enriched pathways and functions in the consensus DAPs, and **D.** Enriched motifs in the DAPs.

**Supplemental Table 3. Protein-protein interactions underlying *ZNF180*-regulome. A.** List of protein-protein interactions from STRING database underlying consensus DEGs and DAPs. **B.** Protein feature table for *ZNF180*-regulome PPI network.

## Notes

### Competing Interest Statement

The authors have declared no competing interest.

