## Supplemental Material for "*ZNF180* modulates tumor intrinsic immunotherapy resistance in melanoma through driving plasticity"

**SUPPLEMENTAL FIGURE**

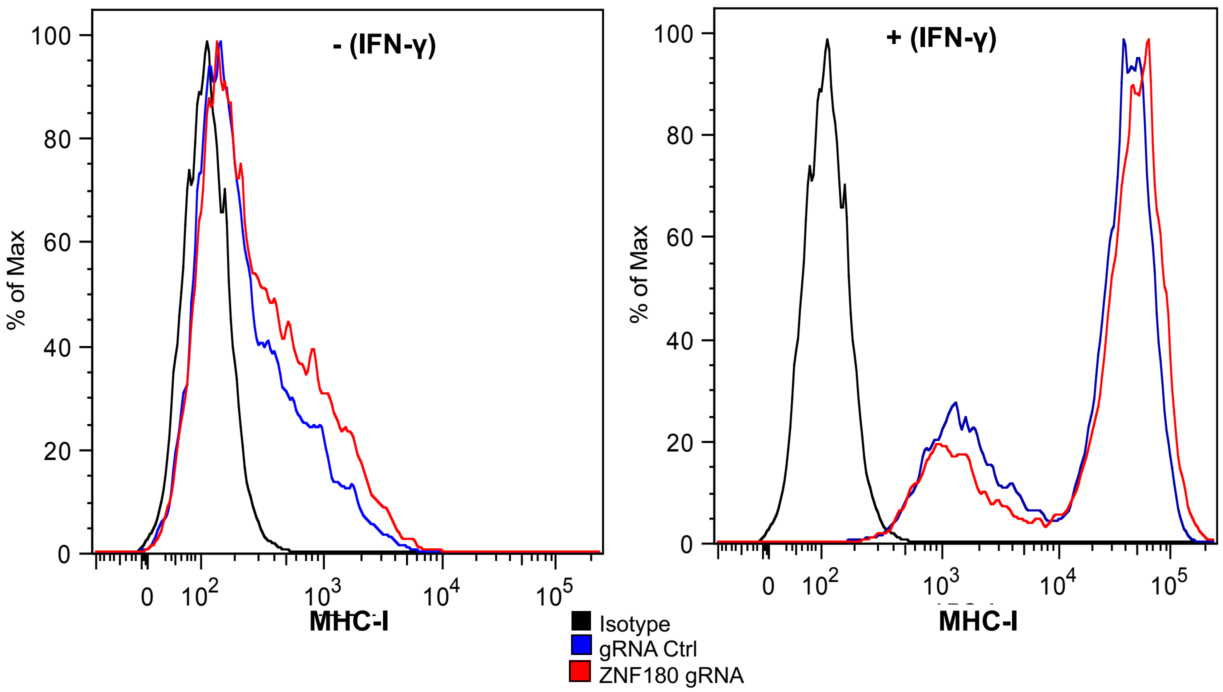


**Supplemental Figure 1.** Raw fluorescence intensity measurements of MHC-I in B16-F12 cells in sgRNA transfected cells (red), control gRNA transfected cells (blue) and isotype control (black).

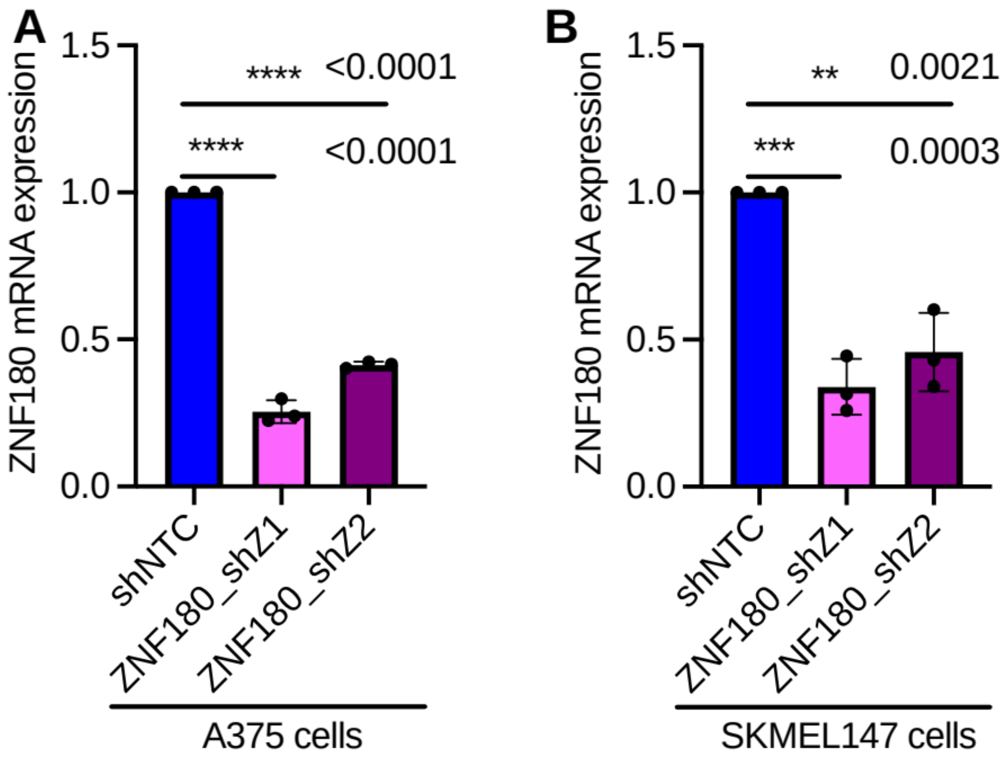


**Supplemental Figure 2. ZNF180 suppressions in A375 (A) and SKMEL147 (B) cells by two shRNA constructs (shZ1, shZ2).**

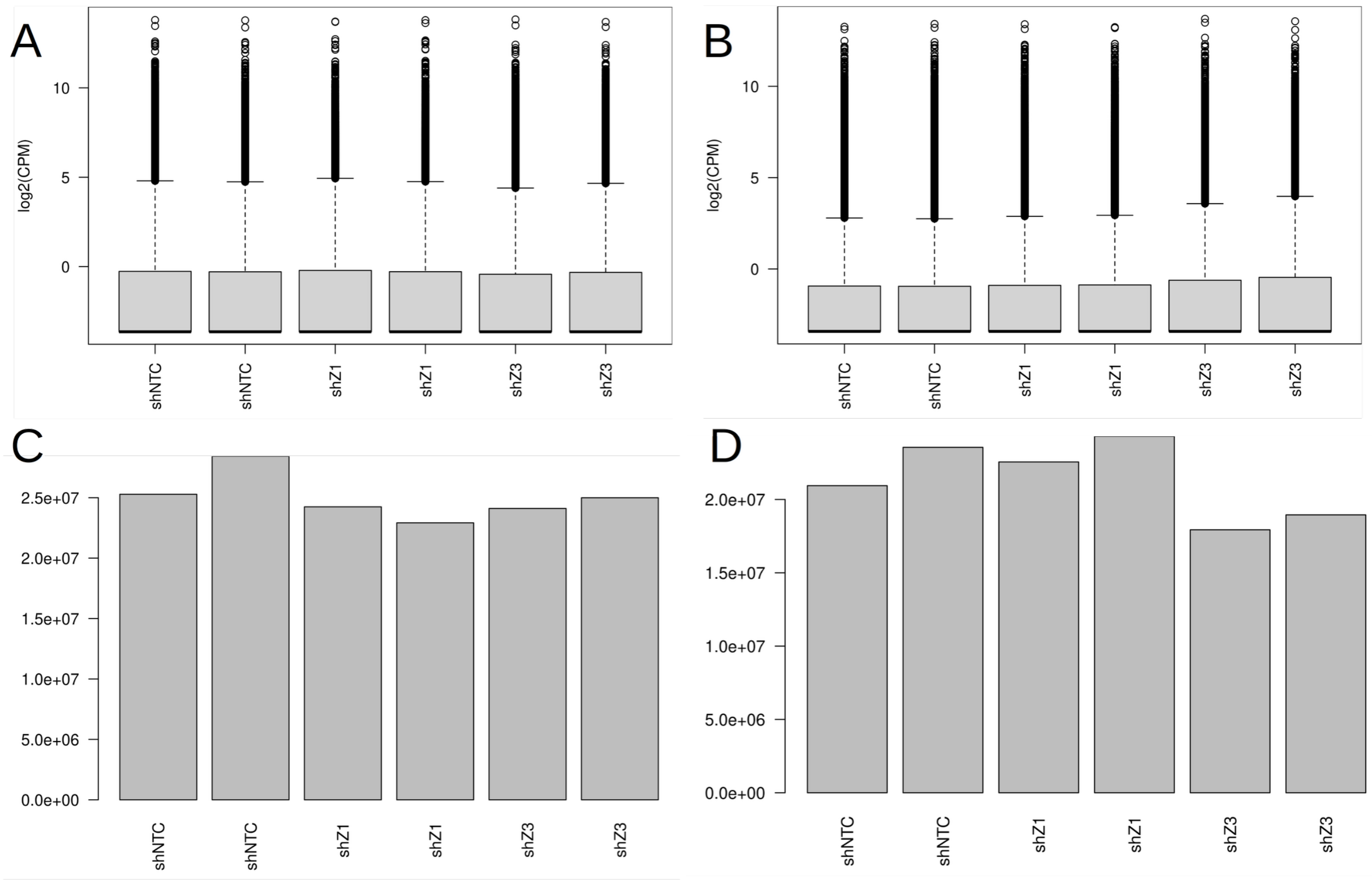


**Supplemental Figure 3. Gene expressions by RNA-sequencing. A, B.** log2(CPM) expressions on different samples from A375 (**A**) and SKMEL147 (**B**) cells. **C, D**. Barplots to show library read depths on different samples from A375 (**C**) and SKMEL147 (**D**) cells.

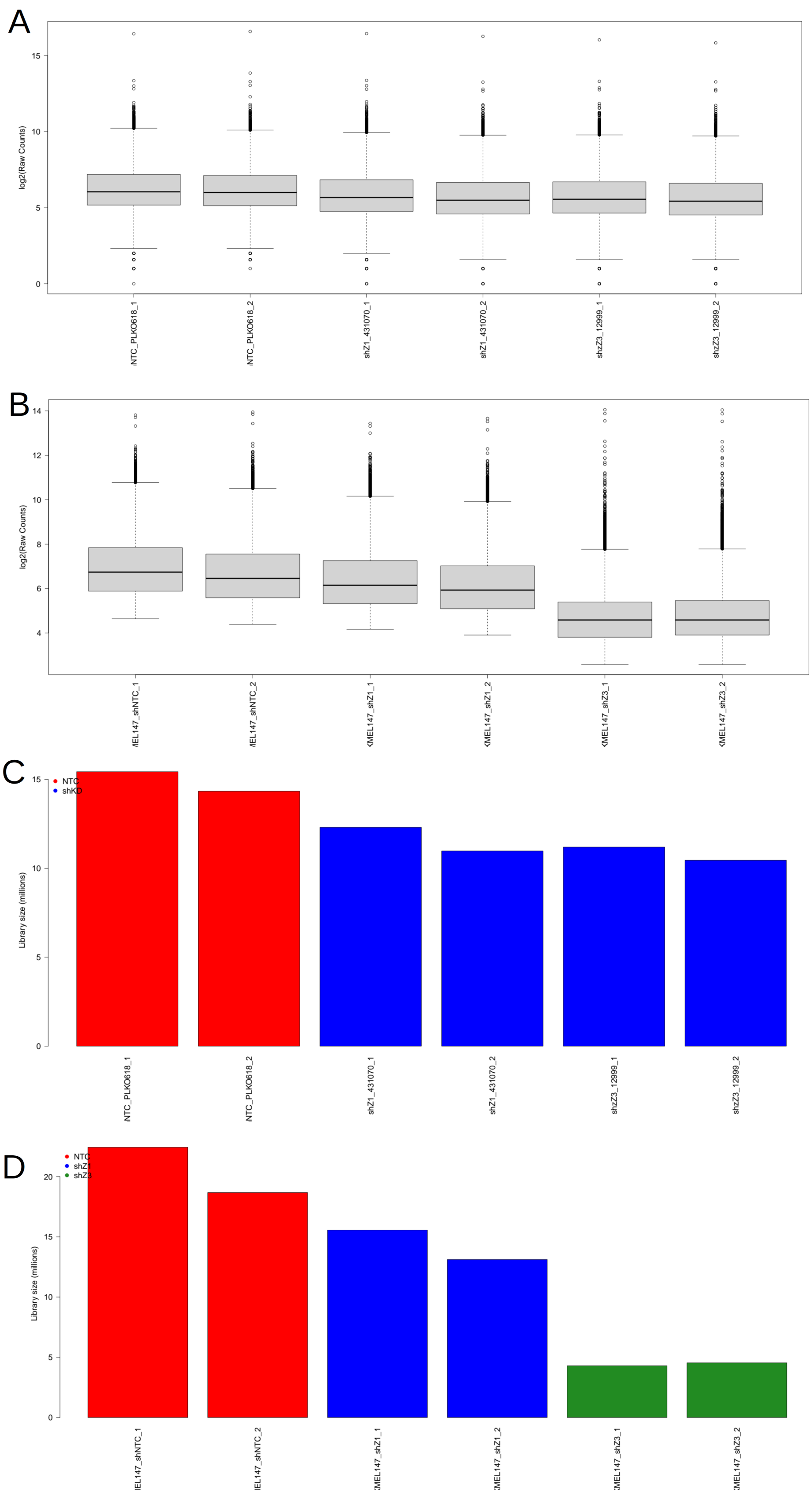
A
log2(Raw Counts)
NTC_PLKO618_1
NTC_PLKO618_2
shZ1_431070_1
shZ1_431070_2
shZZ3_12999_1
shZZ3_12999_2
B
log2(Raw Counts)
C
Library size (millions)
NTC
shKD
NTC_PLKO618_1
NTC_PLKO618_2
shZ1_431070_1
shZ1_431070_2
shZZ3_12999_1
shZZ3_12999_2
D
Library size (millions)
NTC

**Supplemental Figure 4. A, B. Boxplots of samplewise raw read counts from ATAC-sequencing of A375 (A) and SKMEL147 (B) cells. C, D. Barplots of samplewise read depths from ATAC-sequencing of A375 (C) and SKMEL147 (D).**

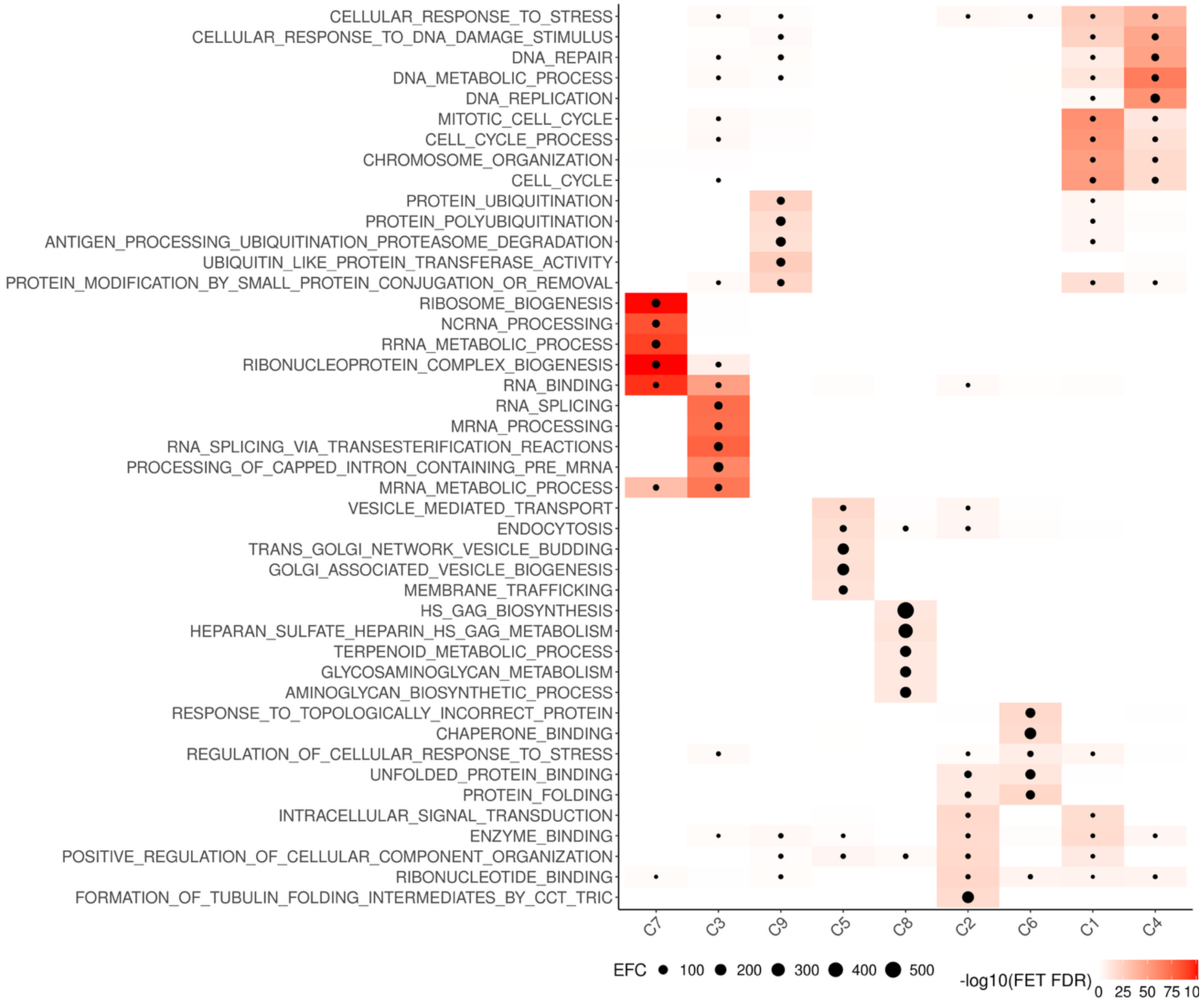


**Supplemental Figure 5. Top 5 most enriched MsigDB pathways and functions in ZNF180-regulome modules.**

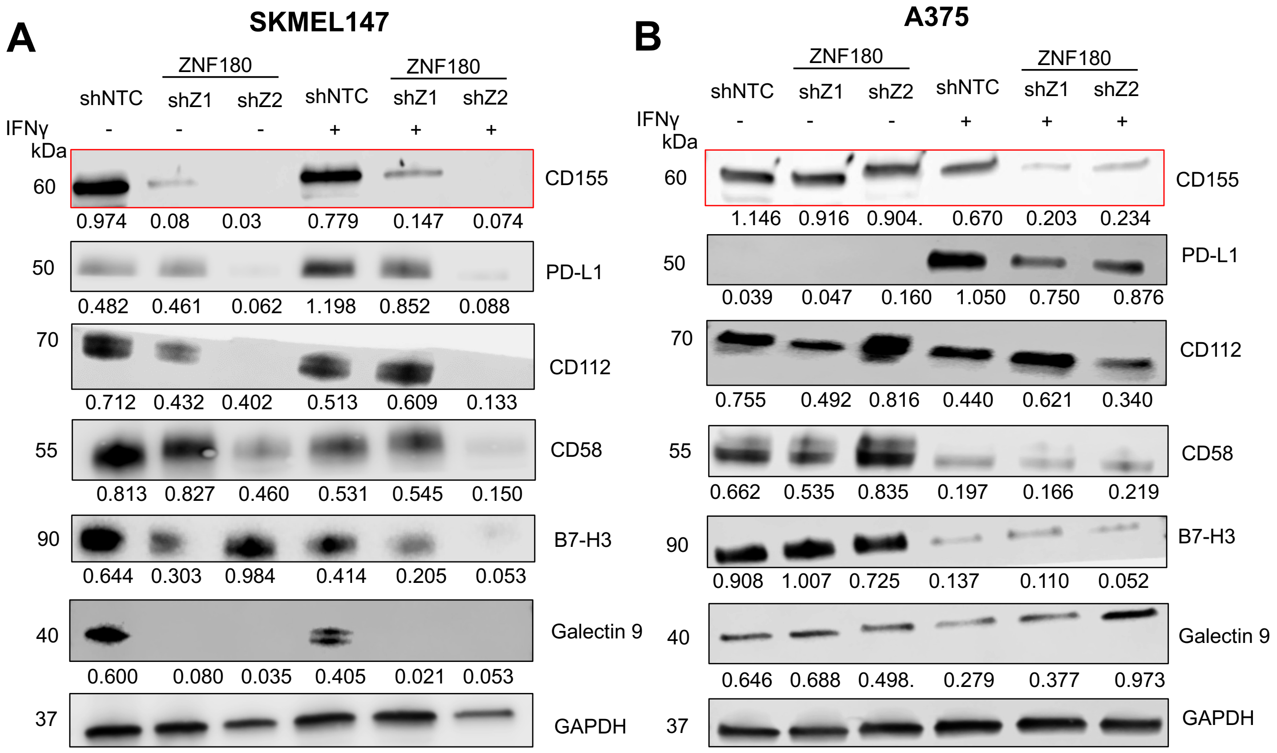


**Supplemental Figure 6. Protein expression of various ligands for T-cell immune checkpoints in ZNF180-silenced (A) SKMEL147 and (B) A375 human melanoma cells.** *ZNF180* silencing resulted in robust reduction of CD155 in both cells, as shown by Western Blot (WB). Cells with IFN-γ+ were treated with 5 ng/ml IFN-γ for 36 hours. GAPDH was used as a loading control as shown on the bottom.

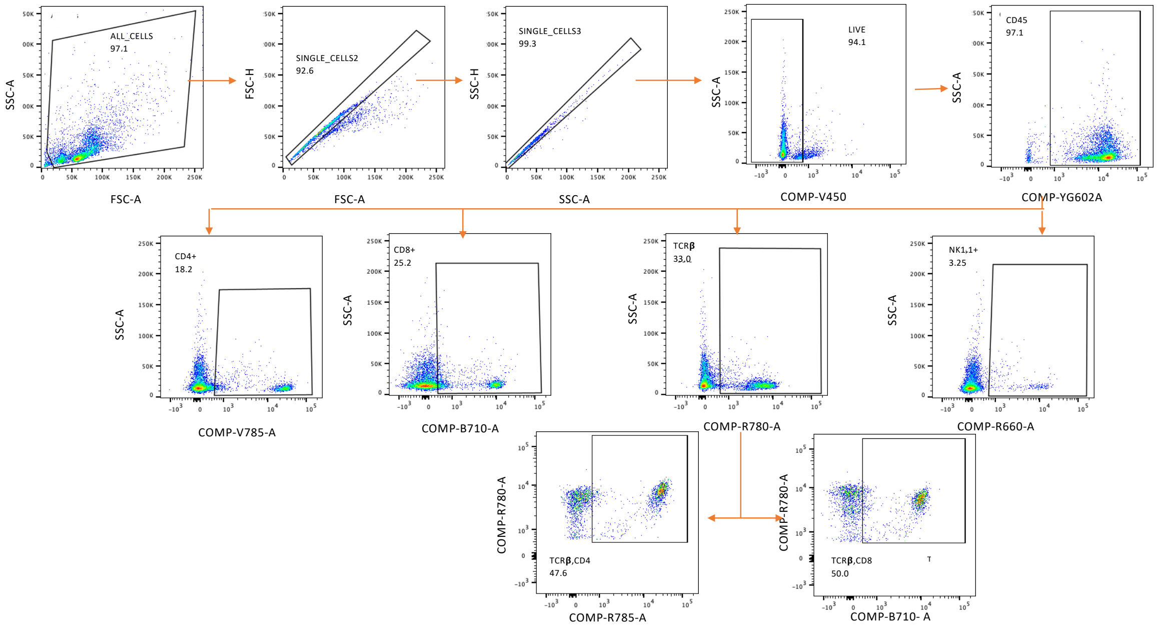


**Supplemental Figure 7. Flow cytometric analysis of tumor-infiltrating lymphocytes (TILs).** Representative gating strategy showing immune subsets isolated from tumor tissues. Live single cells were first gated for $CD45^+$ cells, followed by analysis of $CD4^+$ T cells, $CD8^+$ T cells, $TCR\beta^+$ T cells, and NK1.1$^+$ natural killer cells. Within the $TCR\beta^+$ population, proportions of $TCR\beta^+CD4^+$ helper T cells and $TCR\beta^+CD8^+$ cytotoxic T cells were quantified. Percentages shown in each gate indicate the fraction of positive cells within the parent population.

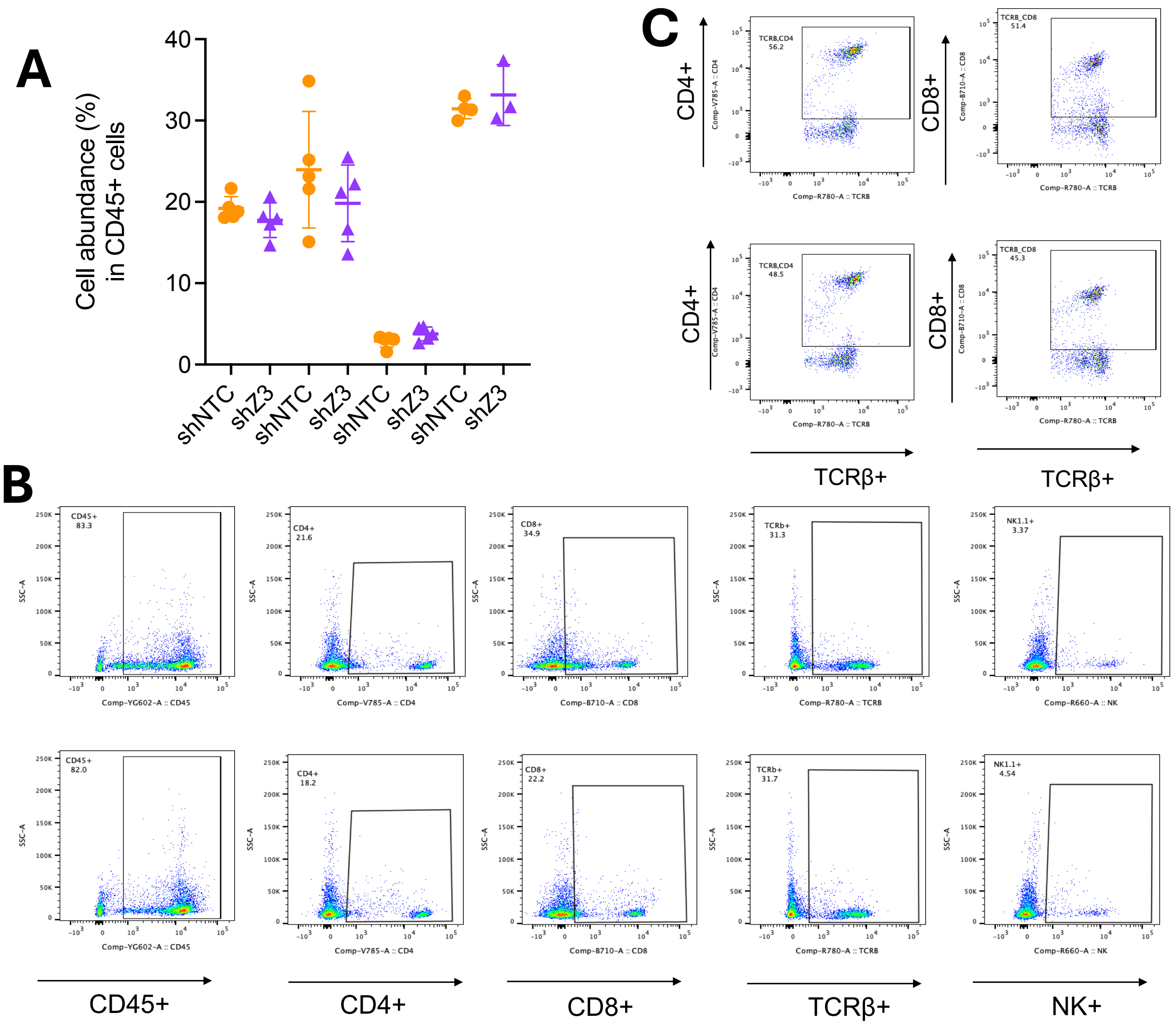


**Supplemental Figure 8. ZNF180 silencing does not alter systemic lymphoid composition in the spleen. A.** Quantification of immune cell subsets in spleens from tumor-bearing mice injected with Yumm1.7 shNTC or shZ3 cells. The frequency of $CD4^{+}$, $CD8^{+}$, $NK1.1^{+}$, $TCR\beta^{+}$, $CD4^{+}TCR\beta^{+}$, and $CD8^{+}TCR\beta^{+}$ populations among splenic $CD45^{+}$ cells was measured by flow cytometry. No statistically significant differences were observed between groups. Data are shown as mean ± SEM. **B, C.** Representative flow cytometry plots showing gating strategies for major immune subsets (**B**) and $TCR\beta^{+}$ CD4+ and CD8+ T-cells (**C**) in the spleens of mice bearing shNTC (top row) and shZ3 (bottom row) tumors.

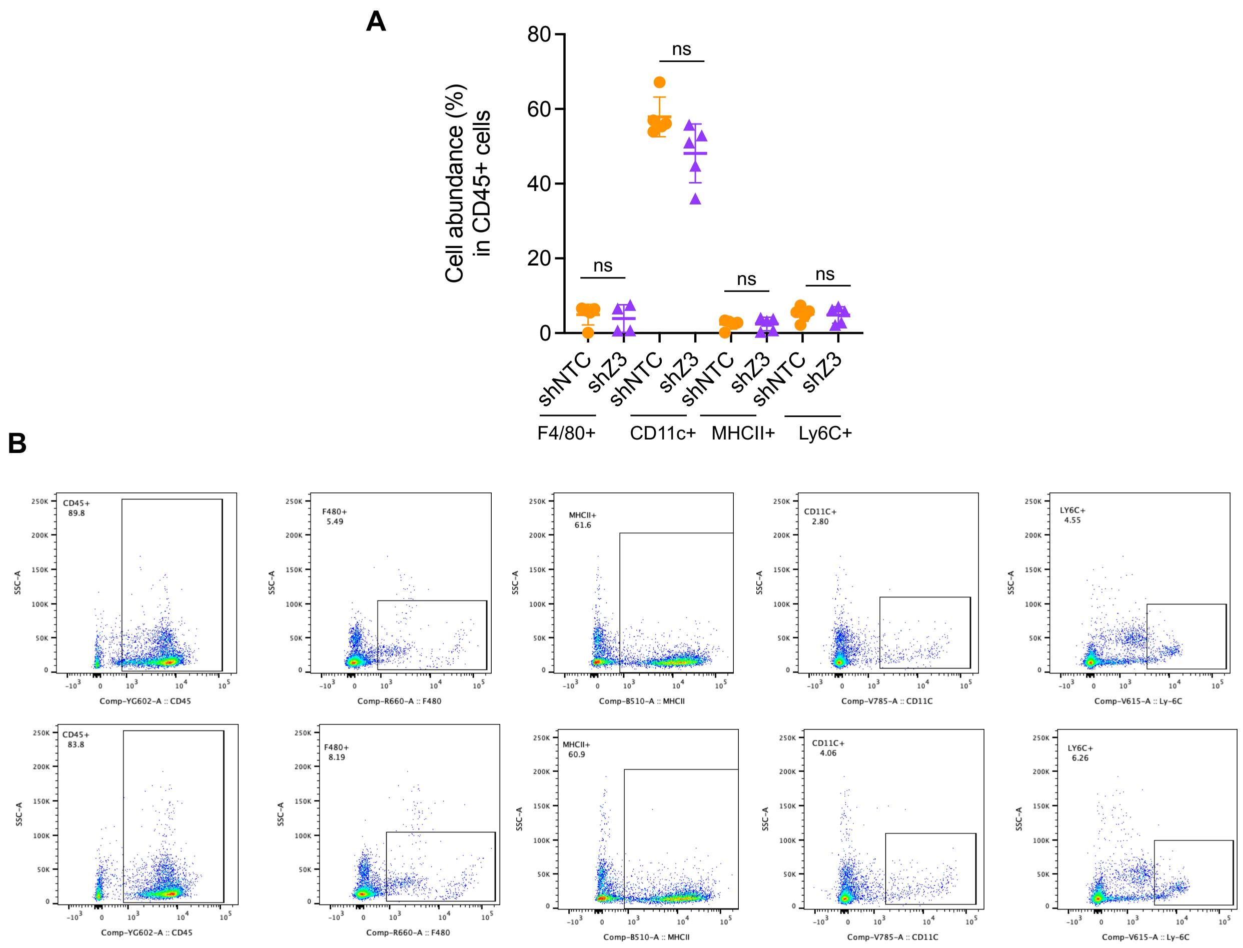


**Supplemental Figure 9. ZNF180 silencing does not alter systemic myeloid composition in the spleen. A.** Quantification of myeloid immune cell subsets among CD45$^{+}$ splenocytes from mice bearing shNTC or shZ3 Yumm1.7 tumors. Frequencies of F4/80$^{+}$ macrophages, CD11c$^{+}$ dendritic cells, MHCII$^{+}$ antigen-presenting cells, and Ly6C$^{+}$ monocytes were assessed by flow cytometry. No statistically significant differences were observed between groups. Data are shown as mean ± SEM. **B.** Representative flow cytometry dot plots illustrating gating for each population in spleens from shNTC (top row) and shZ3 (bottom row) tumor-bearing mice.

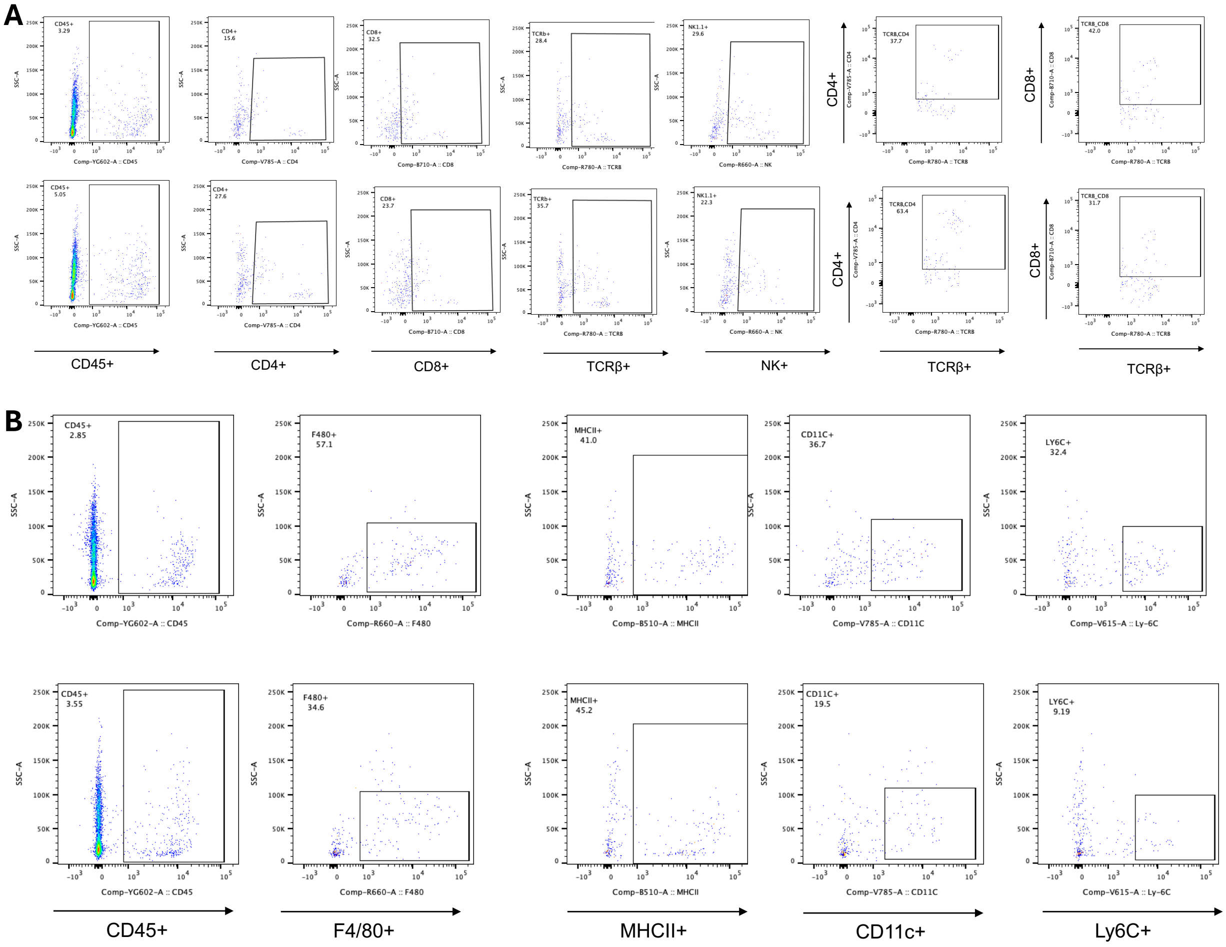


**Supplemental Figure 10. A.** Representative flow cytometry gating strategy and dot plots for immune profiling in Yumm1.7 tumors for lymphoids (**A**) and myeloids (**B**).

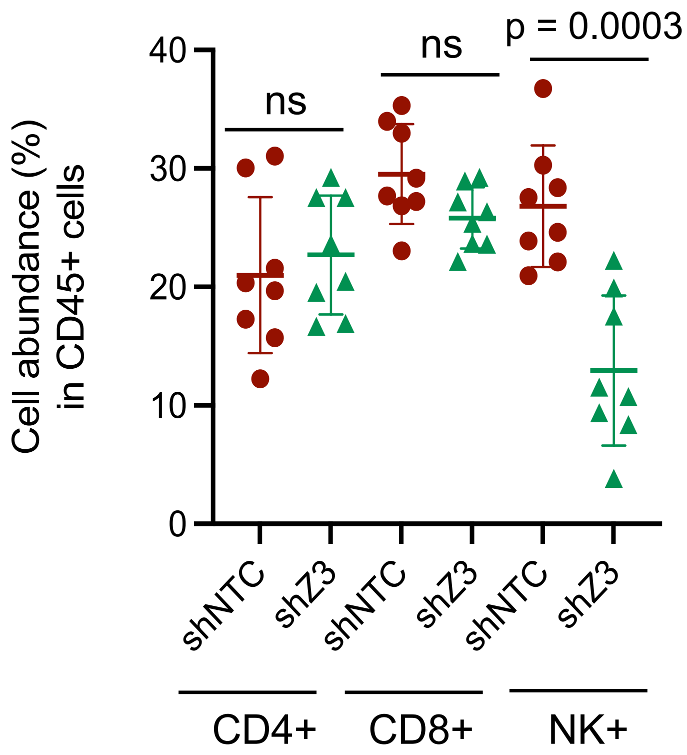


**Supplemental Figure 11.** Flow cytometric quantification of $CD4^+$ T cells, $CD8^+$ T cells, and NK cells ($CD3^-NK1.1^+$) among $CD45^+$ cells. *Zfp180* knockdown significantly reduced NK cell abundance ($p = 0.0003$) with no significant change in $CD4^+$ or $CD8^+$ T cells.

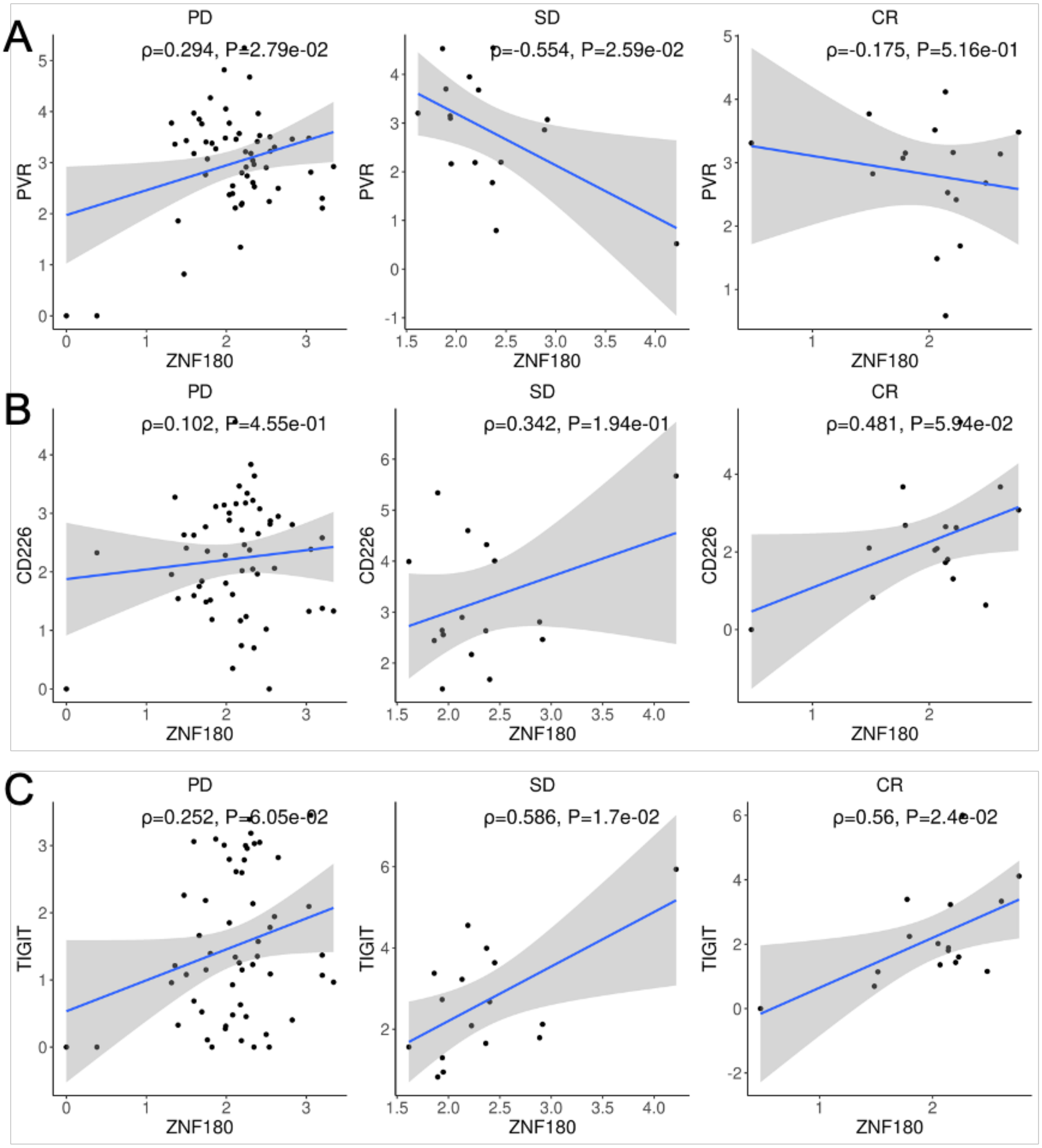


**Supplemental Figure 12. Correlations between *ZNF180* and TIGIT/CD155, DNAM-1/CD155 checkpoints. A.** Correlations with *PVR* (i.e. CD155), **B.** Correlations with *CD226* (i.e. DNAM-1), and **C**. Correlations with TIGIT in C for non-responders (PD) and responders (SD, CR) from Liu *et al.* 2019 study(2).

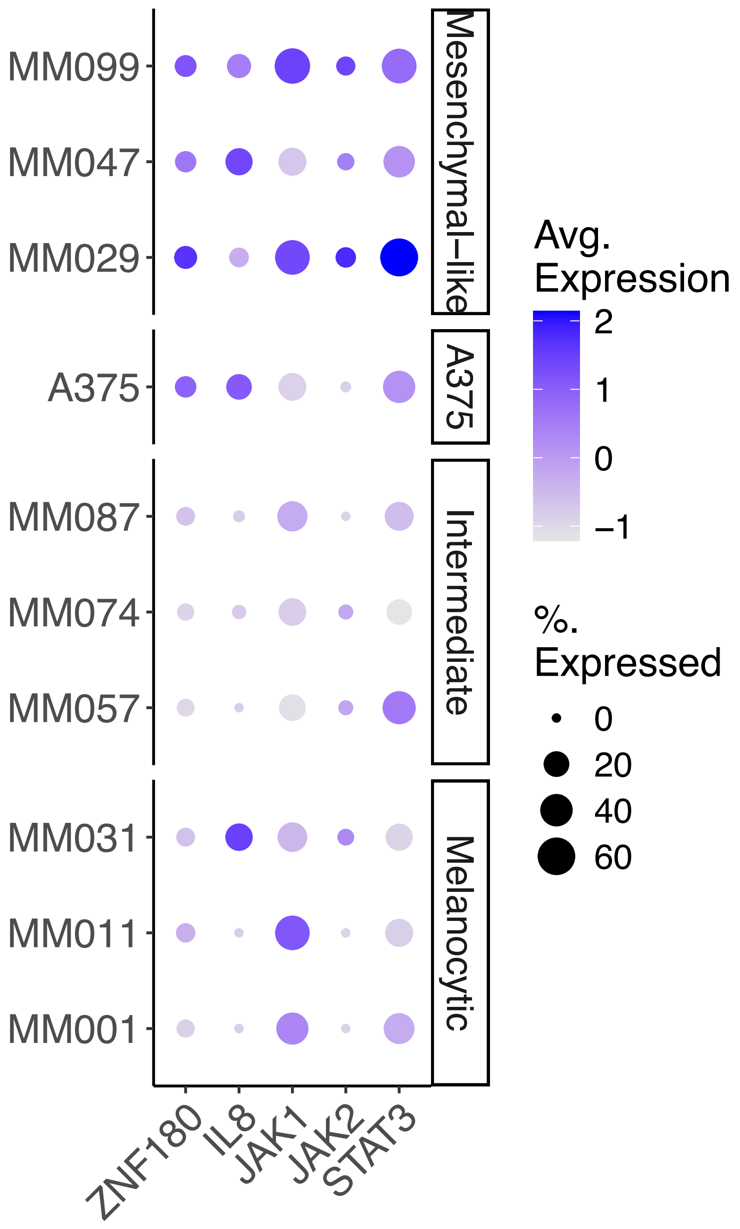


**Supplemental Figure 13.** Dot plot to show co-expression of ZNF180 with IL8 (also known as CXCL8)-JAK/STAT3 pathway in single-cell transcriptomes of patient-derived melanoma cell lines with different subtypes (melanocytic, intermediate and mesenchymal-like) from Wouters *et al.* 2020(1).

**SUPPLEMENTAL TABLE**

**Supplemental table 1. Differentially expressed genes by ZNF180 knock down. A.** List of differentially expressed genes (DEGs) by *ZNF180* knock-down in A375 and SKMEL147 cells, and **B.** Enriched pathways and functions in the consensus DEGs across A375 and SKMEL147 cells.

**Supplemental table 2. Differentially accessible peaks by *ZNF180* knock down. A.** List of differentially accessible peaks (DAPs) by ZNF180 knock-down in A375 and SKMEL147 cells, B. List of consensus DAPs from A375 and SKMEL147 cells, **C.** Enriched pathways and functions in the consensus DAPs, and **D.** Enriched motifs in the DAPs**.**

**Supplemental table 3. Protein-protein interactions underlying *ZNF180*-regulome. A.** List of protein-protein interactions from STRING database underlying consensus DEGs and DAPs. **B.** Protein feature table for *ZNF180*-regulome PPI network.
